# WGCNA+: AI-powered WGCNA for Integration of Multi-Omics Data

**DOI:** 10.64898/2026.09.02.748772

**Authors:** Antonino Zito, Xavier Escriba’ Montagut, Santiago Cano-Muniz, Axel Martinelli, Murodzhon Akhmedov, Ivo Kwee

**Affiliations:** BigOmics Analytics, Via Serafino Balestra 12, Lugano, 6900, Switzerland

## Abstract

**Background:** Weighted Gene Co-expression Network Analysis (WGCNA) is a widely adopted systems biology method to discover gene modules and module-trait associations, mostly from transcriptomics. Designed for a single layer, it cannot jointly analyze multi-omics layers, a consequential limitation in modern biomedical research. WGCNA modules are often hard to interpret, requiring vast follow-up for contextualization. Moreover, no integrated framework exists to visualize condition-specific, cross-omics relationships at module or feature level.

**Results:** To address these limitations, we developed WGCNA+, a novel R package extending WGCNA to multi-omics. WGCNA+ offers key innovations: (i) a unified multi-omics pipeline for per-layer network inference and cross-layer module enrichment; (ii) SVD-accelerated topological overlap matrix calculation that greatly reduces computation time; (iii) a consensus framework identifying modules reproducible across independent datasets/conditions; (iv) LASAGNA, a companion R package for phenotype-conditioned, multi-partite graph visualization of cross-omics relationships; (v) AI-powered annotation and infographics offering immediate biological insight. We tested WGCNA+ across public transcriptomics, proteomics, and miRNA datasets. WGCNA+ detects biologically meaningful modules, cross-omics feature and phenotype correlations, and provides AI-powered interpretation that accelerates research.

**Conclusions:** WGCNA+ addresses existing gaps with a principled, efficient framework for co-expression network analysis across omics. It detects cross-omics regulatory modules and their phenotype association to support basic research, biomarker discovery and pathway analysis. It uniquely offers AI-assisted interpretation and infographics, aiding hypothesis generation. Complementing WGCNA+, LASAGNA is a phenotype-aware multi-partite visualization framework to explore cross-omics relationships. Altogether, these features make WGCNA+ an innovative, powerful tool for clinical and translational research.

**Availability and implementation:** WGCNA+ and LASAGNA are implemented in R language for statistical computing, version≥ 3.5. WGCNA+ and LASAGNA are fully and freely available with no restrictions (https://github.com/bigomics/WGCNAplus; https://github.com/bigomics/lasagna).

## 1. Introduction

Modern biomedical research increasingly relies on the simultaneous profiling of multiple molecular layers, such as transcriptomics, proteomics, metabolomics, and epigenomics. This multi-omics paradigm has fundamentally changed systems biology by enabling the dissection of complex phenotypes through the lens of molecular cross-talk and regulatory cascades that can hardly be captured by any single data layer in isolation. Integrating such data demands analytical frameworks that are not merely capable of concatenating layers but that can rather identify biologically coherent structures across them.

Weighted Gene Co-expression Network Analysis (WGCNA) [Langfelder & Horvath, *BMC Bioinformatics* 2008] is a widely used computational methods in functional genomics analyses. WGCNA, which is almost exclusively adopted in transcriptomics data, identifies modules of co-expressed genes by constructing a weighted adjacency network from pairwise correlations, applies soft-thresholding power transformations to approximate scale-free topology, and clusters genes via hierarchical clustering of the topological overlap matrix. Module eigengenes, representing the first principal component of each module’s expression profile, are used as module representatives that can be correlated with phenotypes to identify biologically meaningful co-expression programs. WGCNA has been applied to numerous studies across neuroscience, oncology and immunology [e.g., Voineagu et al., *Nature*, 2011; Gandal et al., *Science*, 2018; Yang et al., *Nature Communications*, 2014; Hasankhani et al., *Front Immunol*, 2025].

Critically, the standard WGCNA framework has several limitations that restrict its application in multi-omics settings. First, it is conceived for single-omics designs. WGCNA was developed for gene expression data and does not provide a principled mechanism for the joint analysis and integration of distinct molecular layers. For instance, researchers wishing to apply WGCNA to mixed transcriptomics and proteomics data would either need independent runs due to lack of dedicated cross-layer module discovery or enrichment, or limit WGCNA to transcriptomics [e.g., Yang et al., *Int J Onc*, 2019; Johnson et al., *Nat Neuroscience*, 2022]. Second, the current WGCNA has a known computation scalability problem: the full TOM calculation requires O(n³) operations for n features and O(n²) memory. For any dataset comprising tens of thousands features -a near-typical case in modern biomedical research-this imposes a severe computational bottleneck. WGCNA’s blockwise approach partially alleviates this issue by processing features in blocks, however it might result in suboptimal module structure. Third, standard WGCNA enrichment analyses do not account for all omics layers eventually available to inform biological findings. A representative case would be a transcriptomic module significantly correlated with proteomics/metabolomics profiles. WGCNA’s ‘*blockwiseConsensusModules*’ function identifies modules conserved across multiple (transcriptomics) datasets. Yet, no integrated framework exists for integration of distinct omics layers concurrently profiled from the same set of samples, or across experimental cohorts. Furthermore, the detected gene modules in WGCNA may be hard to interpret and often require extensive, time-consuming follow-up analyses for contextualization within biological and biomedical domains. The use of AI to aid biological interpretation and summarization of the results is now widespread. Significant efforts are focused on enhancing existing bioinformatics tools with AI capabilities. The goal is to boost speed, reduce the chance of missing potentially meaningful signals, and rescue neglected profiles. Thus, beyond functions for multi-omics, we have equipped WGCNA+ with tools for AI-powered annotation of results, complemented with infographics. The automated biological interpretation of the results offer immediate biological insight, representing a collaborative platform for biologists to deal with preliminary interpretations and faster decisions about follow-up studies.

Previous efforts have been made to address specific WGCNA limitations and expand its purposes. hdWGCNA (Morabito et al., *Cell Reports Methods* 2023) adapts the framework for high-dimensional single-cell and spatial transcriptomics data through metacell aggregation. Yet, hdWGCNA remains restricted to a single data type. Another extension is CWGCNA (Liu, *NAR Genomics & Bioinformatics* 2024), aimed to include causal inference and multi-omics clustering via canonical correlation analysis. Yet, CWGCNA does not implement a unified co-expression network framework across multiple-omics layers. WGCNA has been applied independently to individual omics layers, but this approach is not suited to capture cross-omics alterations and might also challenge reproducibility. To date, no R package provides a coherent, end-to-end framework for multi-omics WGCNA with integrated cross-layer enrichment, consensus analysis, computational acceleration, and automated reporting.

We present WGCNA+, a novel R package that addresses these limitations through principled multi-omics capabilities [Fig.1]. WGCNA+ significantly expands the scope of WGCNA by adding multi-omics support, computationally efficient topological overlap matrix (TOM) approximation, and rich AI-powered annotation. WGCNA+ introduces: (i) unified per-layer network inference pipeline with cross-omics enrichment; (ii) fast, SVD-based TOM calculation for large-scale analyses; (iii) integrated consensus analysis framework across datasets/conditions; (iv) “LASAGNA” multi-partite graph model, conditioning features on phenotypes; (v) AI-powered module annotation and infographics. WGCNA+ is built from and is fully compatible with WGCNA.

**Figure 1.**
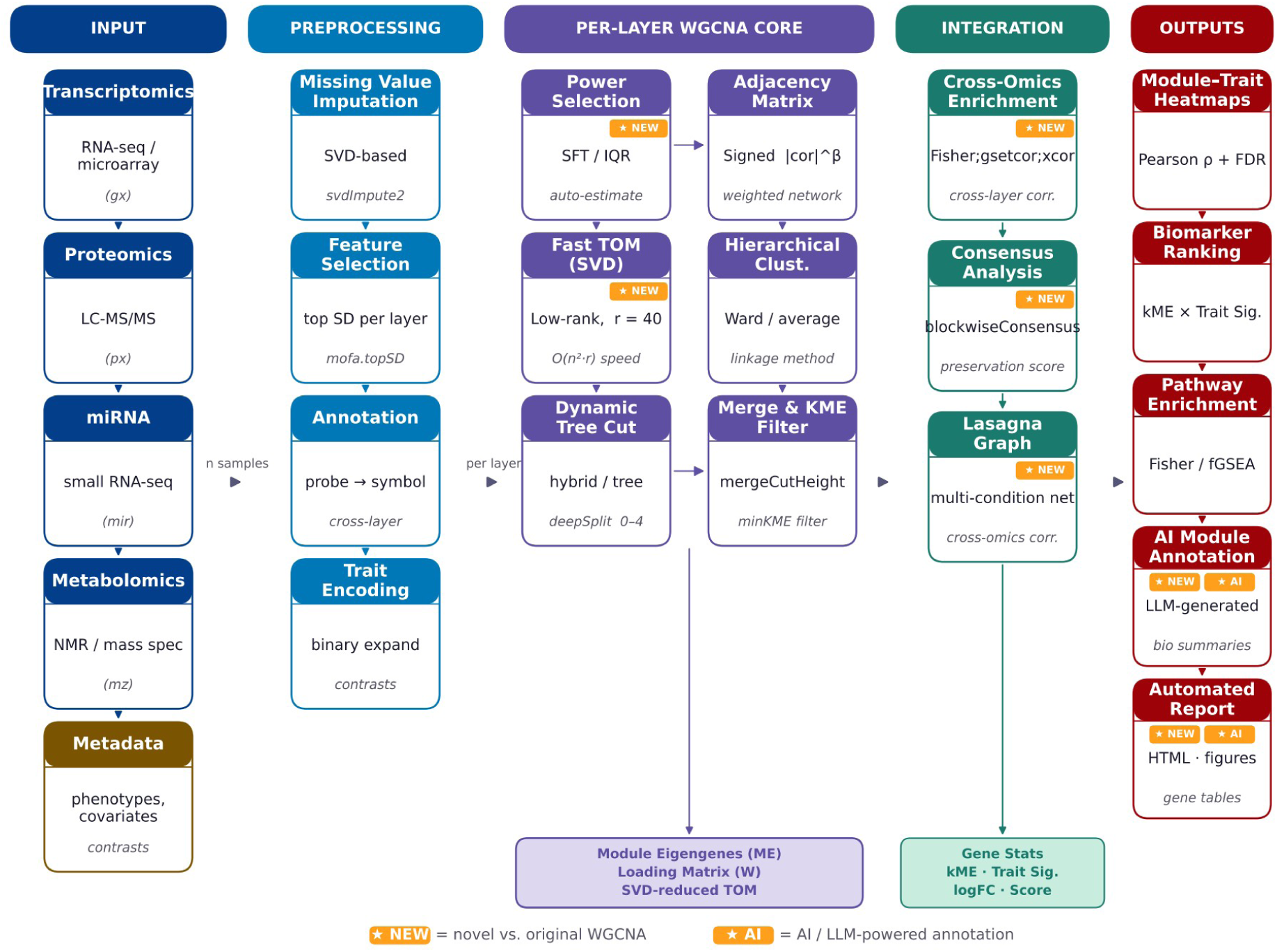
WGCNA+ software overview. Yellow ‘NEW’ and ‘AI’ boxes flag the new functionalities available in WGCNA+. All steps are described in details in the Methods section.

## 2. Methods

### 2.1 Overview of the WGCNA+ framework

WGCNA+ supports four primary analysis modes, all built on a common core pipeline:

1. Single-Omics WGCNA: *computeWGCNA()*. This is the core WGCNA+ function. From a single expression matrix and metadata, it (i) selects the top variable features, (ii) computes a co-expression network using soft-thresholding, (iii) clusters features into modules via hierarchical clustering of a TOM similarity matrix, and (iv) returns module eigengenes, module-trait correlations, and per-feature statistics (module membership, trait significance). The default TOM computation uses a fast SVD low-rank approximation (calcMethod=“fast”) rather than the full WGCNA TOM, making it substantially faster on large datasets when compared to standard WGCNA TOM calculation.
2. Consensus WGCNA: *runConsensusWGCNA().* It tests for modules exhibiting consensus across datasets (e.g., independent cohorts, data types, etc). Each dataset is run through *computeWGCNA()* independently and then WGCNA *blockwiseConsensusModules()* identifies modules consistent across all datasets. Module-trait correlations are computed per-dataset and then summarized into a consensus matrix that only retains statistically significant and directionally concordant associations in all layers.
3. Multi-Omics WGCNA: *computeWGCNA_multiomics()*. It first runs independent WGCNA networks on each data type. After per-layer network construction, a LASAGNA graph model is built on the merged eigengene matrices to capture cross-omics relationships. Enrichment is computed with optional cross-omics correlation in addition to standard Fisher and gene-set-correlation methods.
4. Module Preservation: *runPreservationWGCNA()*. It tests whether modules defined in one dataset are structurally preserved in another. Internally runs consensus WGCNA to establish a reference module set, then calls WGCNA’s *modulePreservation()* to compute Z-summary statistics to assess preservation of each module across datasets.

Critically, all WGCNA+ modes share common sub-routines for feature selection, power estimation, TOM computation, hierarchical clustering, module detection, gene statistics, and enrichment analysis. These sub-routines are modular R functions and are described in detail below.

### 2.2 WGCNA+ input data structures

The primary input to WGCNA+ is a feature expression matrix, where rows are features and columns are samples. Accompanying X is a data frame providing metadata, with samples in rows and traits in columns. Samples names in rows in the metadata data frame need to match the sample names in columns in the feature expression matrix. An optional binary contrasts matrix, with samples in rows and contrasts in columns, can be supplied to add differential comparison groups as additional traits. For multi-omics analysis, a named list of feature x sample matrices, one per data type, is required. Each element in the list would correspond to one molecular data layer. Example of supported layer identifiers include gx (gene expression), px (proteomics), mir (miRNA), mz (metabolomics). Alternatively, a single matrix may be provided in which each feature is prepended with a datatype-specific identifier (e.g., gx:gene1, px:protein1, mz:metabolite1). In this latter case, WGCNA+ automatically identifies the distinct data types and splits the data into separate layers. For consensus analysis across multiple datasets, a named list of features × sample matrices is required. Matrices are aligned by taking common features across all datasets. Sample identifiers need to match across data types (i.e., concurrent multi-omics profiling). Gene set enrichment requires a sparse GMT matrix in genes × gene-sets orientation with binary 0/1 membership values. An optional data frame with a “symbol” column enables probe-to-gene-symbol mapping, applied uniformly to both the expression data and the GMT matrix.

### 2.3 WGCNA+ soft-thresholding power selection

In order to reflect the distinct distributional properties of different molecular data types, power selection is performed independently for each omics layer in multi-omics mode. WGCNA+ implements two automatic power selection methods in addition to manual specification:

**Scale-free topology (SFT) method.** Candidate powers are evaluated over the range {1–10, 12, 14, …, 20, 25, 30, …, 50}. For each power β, a weighted adjacency network is constructed and the degree distribution is fit to a power law. The lowest β achieving a scale-free topology model fit R² ≥ 0.85 is selected, following the original criterions [Zhang and Horvath, *Stat Appl Genet Mol Biol* 2005; Langfelder and Horvath, *BMC Bioinformatics* 2008].

**Interquartile range (IQR) method.** Alternative data-adaptive criterion that selects the power maximizing the IQR of connectivity, favoring networks with well-separated hub and peripheral nodes. This method is more robust in cases where scale-free topology is not achieved (e.g., small sample size). IQR maximizes variation of heights of the dendrogram, promoting separability between groups. It tests a range of power values. For each power: (i) computes signed adjacency matrix; (ii) converts it to TOM similarity; (iii) clusters into a dendrogram; (iv) computes the 25th, 50th, and 75th percentiles of dendrogram heights and their IQR. The power value with the largest IQR is picked. When clusters are well-separated, dendrogram heights are more variable and thus have expectedly higher IQR.

### 2.4 WGCNA+ network construction and fast TOM calculation

**Adjacency matrix (A)**. The adjacency matrix A represents the connection strength between all pairs of genes. A is computed by applying a soft-thresholding power to the pairwise correlation matrix:

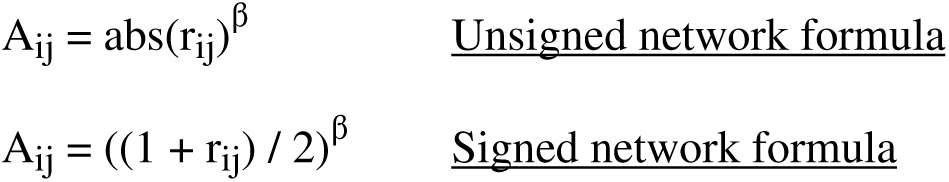

r_ij_ : Pearson correlation between features i and j;

β : Soft-thresholding power value.

Unlike unsigned networks, signed networks preserve the directionality of co-expression relationships. Signed network is the default in WGCNA+.

**Topological Overlap Matrix (TOM).** The TOM generalizes pairwise adjacency by incorporating shared neighborhood information:

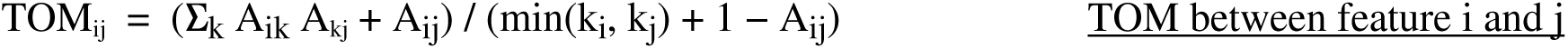

where:

Σ_k_ A_ik_ A_kj_ is the sum of the connection strengths between features i and j to a third overlapping gene k; k_i_ = Σ_j_ A_ij_ is the connectivity of feature i; A_ij_ is the adjacency value representing the connection strength between features i and j. The TOM is more robust to noise than the raw adjacency matrix and improves the separation of modules during clustering.

**Fast TOM via truncated SVD.** Computing the full TOM matrix requires O(n²) memory and O(n³) floating-point operations for n features. For large datasets, this becomes computationally prohibitive. This is a limitation of standard WGCNA. It represents a severe computational bottleneck in modern biomedical studies which often involve large matrices. WGCNA+ addresses this problem by providing an accelerated TOM approximation in the *fastTOMsimilarity()* function. It computes the TOM much faster than the standard WGCNA approach by avoiding the expensive full matrix multiplication step. Specifically, the function exploits the low-rank structure of co-expression matrices. The adjacency matrix A is approximated with truncated (a.k.a. partial) singular value decomposition (SVD) using a low rank component. A is factored using truncated SVD of rank r (default r = 20) via the *irlba()* function from the irlba R package. For robustness, when 20 is higher than half of the number of features in the adjacency matrix, SVD is not computed and a full computation executed. The TOM score is computed for every pair of features setting the self-similarity to 1 so that every feature has perfect topological overlap with itself. Negative values, occasionally arising from low-rank SVD approximation, are set to 0. The TOM is computed from this low-rank approximation rather than from the full adjacency matrix, reducing memory complexity from O(n²) to O(n·r) and computation from O(n³) to O(n²·r). In benchmarking analyses on moderate size datasets of increasing number of features and 18 samples, WGCNA+ reached up to ∼10-fold reduction in TOM computation time compared to WGCNA (Results). WGCNA+ provides three approaches for TOM calculations: ‘fast’ (default), ‘full’ (exact WGCNA TOM), and ‘adjacency’ (uses raw adjacency matrix as TOM). The “blockwise” mode delegates to WGCNA’s blockwiseModules for block-by-block computation on very large datasets.

### 2.5 WGCNA+ hierarchical clustering and module detection

**Feature dissimilarity and clustering.** Pairwise feature dissimilarity is defined as d_ij = 1 − TOM_ij. Features are clustered using agglomerative hierarchical clustering, using the *stats::hclust* function. Supported linkage methods include average (default), complete, single, Ward. This is an advancement in WGCNA+, extending the original WGCNA which only used average linkage. This enhancement would promote more compact and evenly sized modules.

**Dynamic tree cutting.** Module identities are identified by cutting the hierarchical clustering dendrogram using the *dynamicTreeCut* R package. WGCNA+ supports three cutting methods: ‘hybrid’ (default, combines tree topology with TOM distance information), ‘tree’ (topology only), ‘static’ (fixed height). The *deepSplit* parameter controls the sensitivity of the splitting: higher values generate smaller modules, and so a higher total number of defined modules.

**Module merging.** Modules with highly similar eigengenes (as measured by Pearson correlation of module eigengenes across samples) are merged using the *mergeCloseModules* function from the WGCNA package. The merge threshold controls the minimum dissimilarity between module eigengenes required to keep modules separate.

**KME-based module membership check.** Module Membership (kME) is defined as the Pearson correlation between a feature’s expression values and its module eigengene. Features with kME value below 0.3 (default), are removed from their assigned module as considered not representative of the module. Removed features are reassigned to a better-fitting module, or assigned to the ‘grey’ module.

### 2.6 Feature-level statistics

As part of its core computation, WGCNA+ computes the following statistics and metrics:

- **Module Membership (kME):** Pearson correlation of feature expression with each module eigengene, and p-values.
- **Trait Significance (TS):** Pearson correlation of feature expression with each phenotypic trait (continuous or binary-encoded), and p-values.
- **Module-Trait Correlation:** Pearson correlation of each module eigengene with each trait, summarizing module-level phenotypic associations.
- **Differential expression:** for binary traits, limma-based log2-fold changes and p-values are computed for each feature, enabling integration of differential expression results with module membership scores.
- **Compound significance score:** it is defined as the product between kME and TS. It provides a composite measure to identify features that are both strongly module-associated and trait-relevant. This is particularly useful for biomarker prioritization.

### 2.7 WGCNA+ core analysis: *computeWGCNA()*

The central WGCNA+ computation pipeline is implemented in the *computeWGCNA()* function. The following parameters are used per default: soft-thresholding power (power) 12, minimum WGCNA module size (minModuleSize) 20, minimum module membership (minKME) 0.3, threshold for merging similar modules based on the dissimilarity of their eigengenes (mergeCutHeight) 0.15, degree of stringency on how finely the hierarchical clustering tree is split into modules (deepSplit) 2, maximum number of genes (ngenes) to use 2000, tree cut height threshold (treeCut) 0.99, network type ‘signed’. Prior to WGCNA analysis, missing values, if any, are imputed via SVD and duplicated features removed. Features with very low sd (< 0.1 * mean(sd)) are removed and top ‘ngenes’ (by sd) retained. Module detection is achieved using *computeModules(). ComputeModules()* returns same object as standard *WGCNA::blockwiseModules* but is much more computationally efficient and allows different clustering linkage methods (average, ward, complete). A key option in *computeModules* is the ‘calcMethod’ parameter, which specifies the method used for TOM calculation. If calcMethod is “blockwise”, the standard *WGCNA::blockwiseModules()* function is run for network construction and module detection and results returned. If TOM is not provided, the adjacency matrix is calculated using the standard *WGCNA::adjacency()* function. If calcMethod is “adjacency”, TOM calculation is skipped and the adjacency matrix used. If calcMethod is “fast”, *fastTOMsimilarity()* function is used which implements our novel irlba-based low-rank SVD approximation (see paragraph 2.4); if calcMethod is “full”, the standard, computationally inefficient *WGCNA::TOMsimilarity* function is used to compute the TOM from the adjacency matrix. Following TOM calculation, the dissimilarity matrix is generated as 1 – TOM. Hierarchical clustering of features is then performed on the dissimilarity matrix using one of the methods above. The hierarchical clustering results are then subject to adaptive branch pruning using the *dynamicTreeCut::cutreeDynamic* function if the cutting method (cutMethod parameter) is one of “tree” or “hybrid”. If the cutMethod parameter is “static”, the *stats::cutree* function is used to subset the dendrogram into distinct groups. The number of groups “labels” resulting from the dendrogram splitting operation correspond to the number of WGCNA modules. Module colors are then inferred from these labels using the *WGCNA::labels2colors()* function. Module eigengenes are then inferred using the *WGCNA::moduleEigengenes()* function. Pruning based on minKME parameter is then performed to remove, if any, features exhibiting low Pearson correlation with their assigned module’s eigengene. Any feature with KME below the threshold is reassigned to grey module (the “junk” module for unassigned features). To avoid redundancy, highly similar modules are merged using the *mergeCloseModules()* function. Specifically, pairwise Pearson’s correlation values between all module eigengenes are computed and modules are clustered using the average linkage method. Modules exhibiting correlation > 0.85 are merged. Grey module, if present, is not merged with other modules. Modules containing less than *minModuleSize* (default 20) features, are assigned as grey module and effectively ignored in the analysis.

Once *computeModules()* has completed, multi-class categorical variables, if any, are converted into binary strings. To further improve computational efficiency, the full TOM matrix (squared; features x features) is reduced into a low-rank approximation by running *irlba::irlba()*. For each module, the eigen vectors are then calculated; the expression matrix for features in the module is extracted, scaled and centered, and used to compute the first left singular vector via *irlba::irlba()*, corresponding to the per-feature loading onto the module’s first principal component. A loading matrix containing features x modules is then constructed. Note this is different from the module eigengene (which is the sample-space principal components). Per-feature and per-module statistics are then computed using *computeGeneStats()* function. These (i) include modules × traits Pearson correlation and p-value; (ii) correlation between each feature’s expression vs each module eigengene, providing information on how strong is the membership of each feature to the module; (iii) correlation between each feature’s expression vs each trait, providing information on the feature significance score; (iv) for each binary trait, differential profiling is conducted using limma, providing data on each feature log2FC and p-value; (v) for each continuous trait, Pearson’s correlation between each feature and trait is computed; (vi) build a weighted undirected igraph object from the TOM using *igraph::graph_from_adjacency_matrix,* and for each module’s subgraph, runs PageRank using *igraph::page_rank* to score each gene’s topological importance within its module, and finally normalizes centrality values by mean so values are relative. Once *computeGeneStats()* has completed, the module-trait correlation matrix is calculated using Pearson pairwise correlation coefficients between each module and each available trait.

### 2.8 WGCNA+ multi-omics analysis

The WGCNA+ multi-omics analysis pipeline is carried out by the *computeWGCNA_multiomics* function. It extends single-omics WGCNA to a list of data layers. The following analyses are implemented:

**Step 1: Per-layer WGCNA.** Independent WGCNA analyses are performed for each molecular data layer using *computeWGCNA.* Layer-specific soft-thresholding power values, if not specified, are automatically estimated by the function *pickSoftThreshold()*. Within each layer, features receive a two-letter data type-specific prefix (e.g., ‘gx’ for gene expression, ‘px’ for proteomics, ‘mir’ for miRNA).

**Step 2: Consensus layer construction (optional).** For layers sharing a common feature space -typically transcriptomics (gx) and proteomics (px)-, WGCNA+ can optionally construct WGCNA consensus layers using the *createConsensusLayers()* function. This function aligns the two datasets on shared features, runs standard WGCNA *blockwiseConsensusModules(),* and returns consensus module assignments. Consensus modules are WGCNA modules that are consistent across data types. This is particularly relevant for paired transcriptome-proteome studies where concordant module structures may provide evidence of regulatory modules. WGCNA+ consensus analysis is described in detail in Section 2.10.

**Step 3: Phenotype and gene set augmentation (optional).** Two optional augmentation steps are regulated by parameters *add.pheno* and *add.gsets.* These control whether a binary-encoded phenotype variable and gene set scores, respectively, should be appended as additional phenotype layer and treated as pseudo-features. Gene set scores are calculated using PLAID [Zito et al., *Bioinformatics* 2025].

**Step 4: Cross-omics enrichment.** Module enrichment is computed across all layers using three complementary methods. Cross-referencing between layers (e.g., using proteomic features to inform transcriptomic module enrichment) is possible, enabling to specify which layer should be incorporated into the enrichment computation for each target layer.

**Step 5: LASAGNA multi-layer graph.** A multi-condition graph model is built from the per-layer module features. The LASAGNA multi-layer model is described in detail in Section 2.12.

### 2.9 WGCNA+ statistical tests for module enrichment analysis

WGCNA+ implements multiple complementary statistical methods to test for gene set enrichment. These are implemented in the *run_enrichment_methods()* function. All methods can be applied simultaneously to each WGCNA module.

- **Fisher’s exact test.** The Fisher’s exact test is implemented in the *gset.fisher* function. For each module and each gene set, a 2×2 contingency table is constructed based on gene membership (member vs. non-member) and gene set membership (in gene set vs. not in gene set). Benjamini-Hochberg FDR correction is performed. This method tests over-representation of known biological pathways among module genes and is in principle applicable to all omics layers when feature-to-gene-symbol mappings are available.
- **Gene set–eigengene correlation (gsetcor).** Rather than testing binary module membership, this method computes the Pearson correlation between each module eigengene (ME) and the sample-level gene set scores. Gene set scores are computed using PLAID [Zito et al, *Bioinformatics* 2025]. This approach captures continuous module-pathway relationships that may be missed by the binary Fisher test and is particularly informative for large, heterogeneous gene sets.
- **Cross-omics correlation (xcor).** For cross-layer enrichment, the expression profiles of features from one or more reference layers (specified via xref) are correlated with each module’s eigengene from the target layer. Features from the reference layer with high eigengene correlation (|ρ| > 0.8, by default) are used as an extended feature set for Fisher enrichment. This enables, for example, enrichment of a transcriptomic module using correlated proteomic features, linking molecular layers through their shared phenotypic variation.

Enrichment results from all methods are combined per module and ranked by meta p-values. The top enriched gene sets per module are reported in the output and included in automated reports.

### 2.10 WGCNA+ consensus analysis

The consensus analysis framework within WGCNA+ is implemented by *runConsensusWGCNA()*. It aims to identify modules that are reproducible across multiple independent levels (e.g., datasets, cohorts, phenotype traits). Beyond the default parameters in *computeWGCNA()* function (see paragraph 2.7), the following other parameters are used: min p-value threshold for consensus (*cons.psig*) 0.05, minimum genes for enrichment analyses (*gsea.mingenes*) 20, and number of top genes for enrichment (*gsea.ntop*) 1000. The main steps of the consensus analysis are described below in sequential order:

1. <u>Feature alignment</u>: the datasets are intersected on common features. By default, a maximum of 2000 shared features are allowed. If the number of shared features exceeds 2000, the standard deviation (sd) of each feature across samples in each matrix is computed and values of each feature (one for each dataset) are then multiplied to get a combined sd value informing about features with high variability in all datasets. Up to 2000 top-varying features are selected and matrices subsetted to these features.
2. <u>WGCNA format alignment:</u> by default, matrices contain features in rows and samples in columns. For compatibility with standard WGCNA, each matrix is transposed and converted into a multiData format using *WGCNA::list2multiData()* function.
3. <u>Core WGCNA computation</u>: *computeWGCNA()* performs the core WGCNA analysis per dataset.
4. <u>Identify consensus WGCNA modules</u>: The standard *WGCNA::blockwiseConsensusModules()* is run across all datasets simultaneously to detect consensus modules across datasets by using a consensus TOM. The values of soft-thresholding power from individual layer runs are reused. Each individual layer’s module colors, assigned by *computeWGCNA()*, are remapped to the consensus color scheme. Feature names are attached to each block’s consensus dendrogram.
5. <u>Build consensus matrix and compute module-trait correlation and statistics</u>. Pearson pairwise correlation between consensus modules and traits is computed, resulting into a module-trait correlation matrix for each dataset. A single consensus module-trait matrix is produced by *computeConsensusMatrix().* The consensus matrix includes only modules that are individually significant and directionally consistent across all layers. The consensus value has a ‘sign’ and a ‘magnitude’ component. The magnitude component is calculated via three alternative, optional approaches: ‘min’ (default), which takes the minimum absolute value of each layer’s raw matrix and multiply this by the ‘sign’ of the consensus (most conservative estimate, the default WGCNA consensus approach to capture effects that are at least strong as the weakest dataset); ‘gmean’, which calculates the geometric mean of the absolute value and multiply it by ‘sign’ of consensus; ‘mean’, which is the average value. Modules exhibiting discordant sign across layers or statistically not significant in all layers are removed. Consensus statistics are computed using *computeConsensusGeneStats(),* the consenus-aware equivalent of *computeGeneStats()* (paragraph 2.7). If a GMT file with genesets and pathways is supplied, consensus module enrichment can be tested using *computeConsensusModuleEnrichment().* Gene sets and pathways containing fewer than 3 features are removed. This stringency can be adjusted by the user (*min.genes* parameter). Statistical enrichment is conducted using *run_enrichment_methods()* (paragraph 2.9).
6. <u>LLM-powered biological description</u>: for each consensus module, the *describeModules()* function generates automatic, AI-written summaries. The function starts by pulling, for each consensus module, the top 50 correlated genes, top enriched gene sets, and top correlated traits using *getTopTables()*. For each module, a detailed summary is by default requested to the LLM model to gather information on module biological function, roles of top enriched genesets belonging to the module, and possible relationship with phenotypes. This request is passed to the LLM via a static prompt. If a valid AI model is passed and exists, the prompt is sent to the LLM using *ai.ask()*. If no AI model is recognized or available, it falls back to a templated summary. The function has successfully been tested with groq, grok, and open-ai. While the architecture to use LLM is entirely provided by WGCNA+, the purchase and use of the API key is a user’s responsibility. A description of the WGCNA+ LLM capabilities is provided in paragraph 2.13.

### 2.11 WGCNA+ preservation analysis

Preservation analysis in WGCNA+ is implemented by *runPreservationWGCNA(),* a variant of *runConsensusWGCNA().* In addition to finding consensus modules, it tests whether WGCNA+ modules in a reference dataset are structurally preserved and statistically reproducible in another dataset [Langfelder et al., *PLOS Computational Biology* 2011]. To align with current standards, it uses WGCNA’s modulePreservation Z-summary. Default parameters include ‘power’ 12, ‘ngenes’ 2000, ‘minModuleSize’ 20, ‘deepSplit’ 2. The function first builds consensus network using *runConsensusWGCNA()* on all input layers. No consensus stats and module enrichment are tested. It runs *WGCNA::modulePreservation()* with reference network’s module colors, signed network type and 10 permutations. It tests whether the reference modules’ density and connectivity are preserved in other datasets. Preservation is quantified using Z-summary, which combines multiple preservation statistics (density, connectivity, separability). Modules with Z-summary > 10 are considered strongly preserved; modules with Z-summary < 2 are non-preserved; modules with Z-summary between 2 and 10 are weakly/moderately preserved. Once preservation analyses is completed, the module eigengenes per layer are recomputed in each dataset using the color coding of the reference set. Aligning color coding of modules across layers allows comparisons between correlation matrices. Module-trait correlation matrices are then computed per layer. Finally, feature statistics and geneset/pathway enrichment are computed in the reference layer using WGCNA+ *computeGeneStats()* and *computeModuleEnrichment()*.

### 2.12 LASAGNA Graph Model

L.A.S.A.G.N.A (<u>L</u>ayered <u>A</u>pproach to <u>S</u>imultaneous <u>A</u>nalysis of <u>G</u>enomic and <u>N</u>etwork <u>A</u>ssociations) is a novel, self-contained R package. It complements WGCNA+. LASAGNA has primarily been conceived for solving and visualizing correlation-based multi-partite (multi-layer) multi-omics data graphs. In the LASAGNA object, each layer is an omics type, such as transcriptomics (‘gx’), proteomics (’px’), metabolomics (‘mx’), miRNA (‘mir’) etc. The algorithm and its plotting capabilities are described below.

1. The LASAGNA core function is *create_model(),* which create LASAGNA multi-layer graph model. It accepts a list of feature matrices (X), one per omics layer, and a sample metadata table. It constructs a single merged feature matrix with layer-prefixed row names (e.g., “gx:TP53”, “px:TP53”, “mx:Glucose”). Optionally, each layer is filtered to its top (1000 by default) most variable features. Sample metadata are expanded into a numeric design matrix or alternately, converted from a label matrix into signed contrasts. The phenotype information will represent a ‘pheno’ layer in the graph. All layers are merged into a unified multi-omics matrix. A global feature-feature Pearson correlation matrix is computed. Data can be optionally conditioned by phenotype. In this case, Pearson correlation between features and phenotype(s) is computed and the outer product of the per-node maximum absolute phenotype correlations is calculated. The feature-feature correlation coefficients are then multiplied by the outer product. This approach allows to emphasize edges connecting phenotype-relevant features relative to general correlation noise, therefore rendering the graph structurally biased toward phenotype-relevant connections. At this stage, all cross-layers connections can be optionally shown, in which case edges between non-adjacent layers will be also displayed as well as intra-layer correlation edges. By default, the edges are topologically restricted, such that only features mapped to adjacent layers are connected. Edges connecting non-adjacent layers are masked (assigned value of 0). Similarly, intra-layer edges can be optionally masked. In order to ameliorate the quality of the graph and emphasize potentially more relevant features, inter-layer associations are reduced to the top set (by default 20) of most correlated edges per node. These represent the top cross-omics correlated features. Restriction is applied to intra-layer nodes. Finally, an igraph object is created using the *igraph::graph_from_adjacency_matrix()* function. Each data type becomes a named partition of nodes in the graph. In the object, each edge carries a rho attribute and a connection_type label such as “gx->px” for inter-layer or “gx” for intra-layer edges.
2. The LASAGNA igraph object is solved for a single phenotype by the *solve()* function. This function takes the output from the *create_model()* function. It starts by calculating the node statistics: for each feature in the merged multi-omics matrix, the correlation with the phenotype vector is computed and log2FC between phenotypes also calculated. Both correlation and log2FC values are stored as vertex attributes in the igraph object. The user can optionally show log2FC or correlation values at each node in the graph. The geometric mean between the phenotype correlation (or log2FC) of each edge’s two endpoints (nodes) is calculated and multiplied by the original feature-feature correlation. This approach sets the new edge weights: edges between two features that are both strongly phenotype-associated get boosted. Conversely, an edge between two phenotype-irrelevant features gets shrunk toward low values. Through this step the graph becomes condition-specific. Optionally, shortest-path weighting can be enabled, which finds the shortest path through the graph using the reciprocal of each node weight as path cost, such that strong edges are cheap while weak edges are expensive. Each edge is scored by the weakest link along its best path. To generate a clean and high-quality graph, pruning is performed: (i) edges are filtered to include only those with weight above a pre-imposed correlation value; (ii) per each connection type (e.g., ‘gx→px’), only the top 100 edges by absolute weight are retained while all the others are zeroed; (iii) zeroed edges and disconnected nodes are by default removed from the graph.
3. The LASAGNA igraph object is solved for multiple phenotypes by the *multisolve()* function. This function takes the output from *create_model().* For each available phenotype, it first runs *solve()*. No pruning is performed such that the same full vertex set is retained across every phenotype’s solved graph. All resulting per-phenotype graphs are combined into a single feature x trait matrix, which is squared and averaged per feature. Each phenotype-specific adjacency matrix is retrieved from each graph and squared. It then calculates the root mean square per edges across all phenotypes. This approach is sensitive to both phenotype-specific signals and edges consistently robust across multiple phenotypes. Finally, the consensus graph is rebuilt using the multi-pheno data. Specifically, the sign of each weight is recovered from the base correlation graph and the signed weight multiplied by root mean square calculated above. Edges with weight below a pre-imposed threshold are removed. In the rebuilt multi-trait consensus graph, nodes and edges are robustly associated across multiple phenotypes rather than being condition-specific.
4. The LASAGNA igraph object is graphically displayed through a series of advanced plotting functionalities. Multiple visualization functions operate on a solved graph: (i) *plot_multipartite()* draws a static base R plot in either ‘parallel’ or ‘hive’ layout. In the ‘parallel’ layout, layers are displayed as vertical columns, nodes positioned vertically by value, and PageRank-scaled vertex sizes displayed. In the ‘hive’ layout, layers appears as radial axes and nodes populate each axis. Edges are colored differently for positive and negative correlations. In both layouts, nodes’ sizes are relative to centrality and nodes’ colors reflect positive or negative sign; (ii) *plot_visgraph()* produces an interactive visNetwork with physics-based force-directed layout; (iii) *plot_3d()* builds an interactive plotly 3D plot where each omics layer is rendered as a semi-transparent horizontal plane at a different z-level, nodes are scatter-plotted within their plane using 2D coordinates, and inter-layer edges drawn as thin 3D segments colored by sign.

### 2.13 WGCNA+ AI-powered module annotation

WGCNA+ provides AI-powered biological interpretation of results. Some WGCNA+ AI capabilities are described in paragraph 2.10. It follows a detailed description,

1. *create_report()*: this function implements a multi-stage LLM pipeline that turns a WGCNA result object into a detailed narrative report. This prose structure, instructed prior to running the model, facilitates integration into existing or new reports, contrary to typical, bullet points AI-written content. The function first pulls top phenotype-correlated modules using *getTopModules()*. For each module, the *describeModules()* function is called specifying no AI model to generate templated per-module summaries including key features, genesets/pathways, and correlated phenotype (paragraph 2.10). For each module, it generates a summary prose from descriptions, describing underlying biology, correlated phenotypes and mention key features. As multiple AI models might have been used to generate independent module descriptions, this step would smooth potential model disagreements into a more coherent prose. If no AI model is specified or the model is unrecognized/not working, the templated summaries from *describeModules()* are used. A final, higher-level LLM call receives all summaries and generates an integrated scientific-article-style report, connecting modules into biological programs and beyond, including proposing disease-related aspects when possible. Finally, *create_diagram()* is called.
2. *create_diagram()*: this function receives a final LLM-generated report and generates a GraphViz directed graph in DOT format. Connections are built from scratch or constrained by an existing igraph template (e.g., co-expression network, LASAGNA). In the diagram, WGCNA+ modules and phenotypes are connected using LLM, with self-correction and validation. If an existing graph is supplied, this will be used as a template to generate the DOT diagram, constraining connections to it. This approach would reduce possible hallucinations. If no graph is passed, the LLM will generate the graph from scratch by connecting the WGCNA+ modules and relying on external knowledge. Instructions are provided to the model, such as “do not use special characters, subgraphs, hexadecimal color coding”, “use solid lines for positive regulation”. The output is subject to cleanup operations to improve the quality. The generated DOT code is validated and self-repaired iteratively, with *grViz()* and *export_svg()*. The LLM gets re-prompted with any invalid code until a syntactically correct diagram is produced, up to 5 attempts.
3. *create_infographic().* This function takes the whole *create_report()* and appends both report and diagram, if present, to a prompt asking for a graphical abstract in scientific style, publication-ready. It calls *ai.create_image_gemini()*. It aims to generate a rich infographics summarizing the entire report.
4. *create_module_infographic()*. This function is module-specific. It takes the full report object rpt, returned by *create_report()*, and a single WGCNA+ module name and ask *ai.create_image_gemini()* function to generate an infographic summarizing the biological narrative of the module.

### 2.14 WGCNA+ implementation and availability

WGCNA+ and LASAGNA are implemented in R version≥ 3.5 and are freely available on GitHub (https://github.com/bigomics/WGCNAplus; https://github.com/bigomics/lasagna). WGCNA+ can be fully integrated within existing WGCNA workflows. WGCNA+ imports the R packages ‘WGCNA’, ‘irlba’ (SVD), ‘dynamicTreeCut’ (dynamic tree cutting), ’fgsea’ (enrichment), ‘fastcluster’ (fast hierarchical clustering), ‘igraph’ (graph operations), ‘matrixStats’ (fast matrix statistics), ‘Matrix’ (sparse matrix support), ‘ggplot2’ (visualization). Optional dependencies: ‘ComplexHeatmap’, ‘plotly’, ‘uwot’, ‘SuperCell’, ‘DiagrammeR’. The WGCNA+ and LASAGNA vignettes demonstrate complete workflows.

## 3. Results

Among the advances introduced by WGCNA+ is a computationally efficient method for calculating the TOM using truncated SVD. Specifically, the adjacency matrix A is approximated with truncated SVD using a low rank component (Methods). The TOM is computed from this low-rank approximation rather than the full adjacency matrix, substantially reducing computational time. To evaluate the performance, we compared the WGCNA+ implementation with the standard WGCNA TOM calculation using a proteomic dataset of 18 samples [Wolf et al., *Nat Immunol* 2020] at increasing feature space (top 1,500; top 3,500; and all 6,987 features) [Fig. 2].

**Figure 2.**
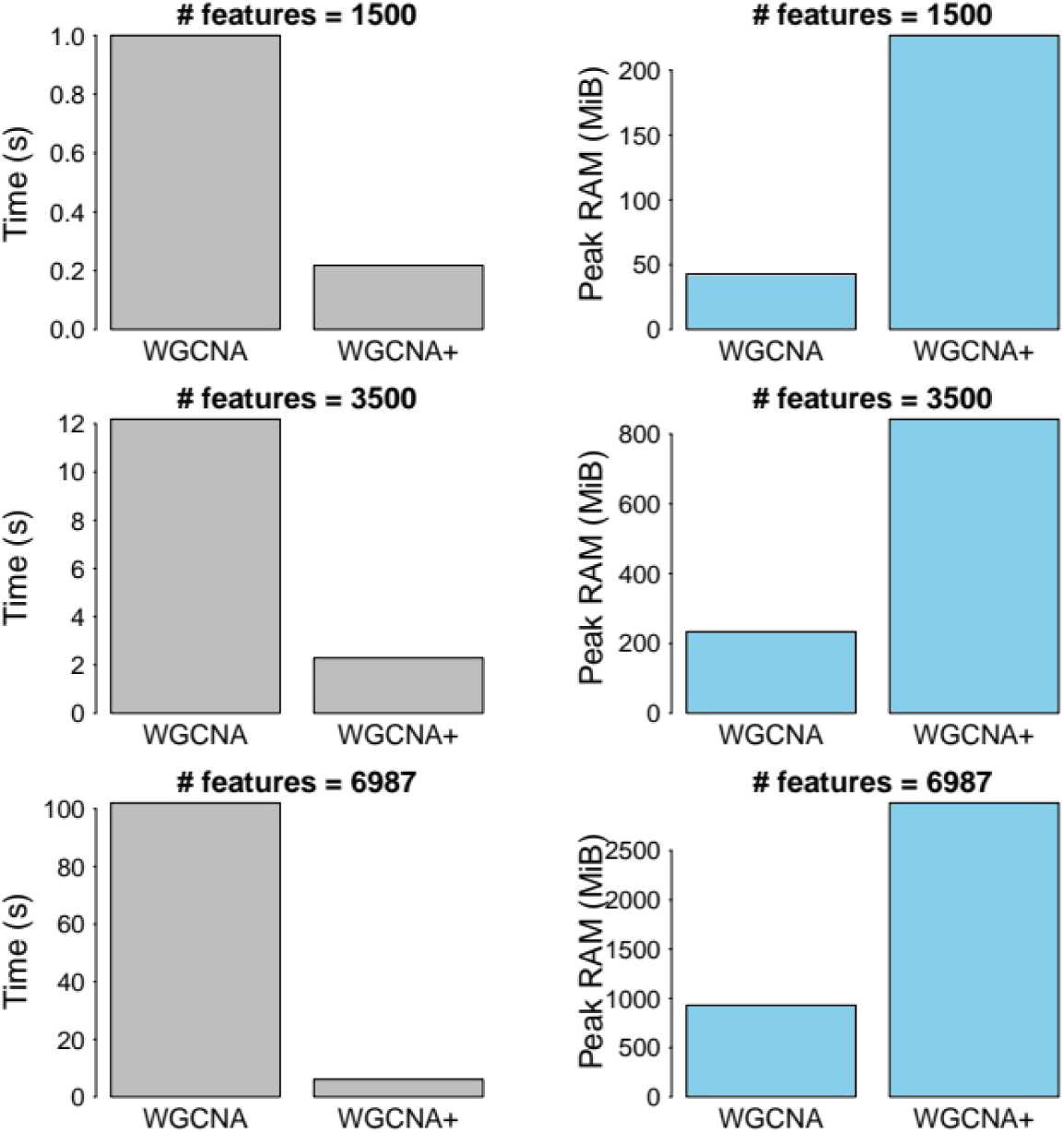
Comparison between WGCNA and WGCNA+ TOM calculation. Runtime vs number of features and peak RAM usage vs. number of features for WGCNA and WGCNA+ TOM calculation step. Comparison conducted using a proteomics dataset [Wolf et al., *Nat Immunol* 2020] containing 6987 features profiled in 18 samples. An increasing number of features (top 1500, 3500, 6987 by standard deviation) was used for testing. Run time (s) was calculated using the *system.time* R function. Peak RAM (MiB) used was calculated using the *peakRAM::peakRAM* R function. The adjancency matrix was first generated using the *WGCNA::adjacency* function. The TOM was calculated from the adjancency matrix using *WGCNA::TOMsimilarity* and *WGCNA+::fastTOMsimilarity* functions.

We demonstrate that the WGCNA+ TOM computation is consistently faster than standard WGCNA, achieving from ∼5 to ∼10-fold increase in calculation speed, albeit with higher RAM usage. The relatively increased RAM usage reaches up to ∼2.5 GB for the full dataset, which remains very well within the capabilities of most modern computing systems used in biomedical research, including standard laptops, and that therefore does not impose a significant problem in present day bioinformatics. On the other hand, the gain in computation speed was excellent, enabling more analyses of large datasets.

We tested WGCNA+ in distinct publicly available datasets, spanning diverse scenarios in terms of size and data types, at both single and multiple omics level [Wolf et al, *Nat Immunol* 2020; https://www.cbioportal.org; https://github.com/edo98811/WGCNA_official_documentation]. Using a proteomics dataset of the dynamics of mRNA translation kinetics and protein turnover in human naive and activated T cells [Wolf et al., *Nat Immunol* 2020], WGCNA+ identified the blue, turquoise and yellow to be the protein modules most significantly associated with the activated immune cell state [Fig.3A; Table S1]. Specifically, blue module is positively correlated with immune cell activation, while the turquoise and yellow modules are negatively correlated with the immune cell activation. Gene set and pathways enrichment analyses, conducted with three independent statistical methods (Methods), revealed that features within the blue module are enriched for multiple drug-related pathways, distinct estradiol-involving signal, and interleukin-related gene sets [Table S2]. By contrast, features mapped into the turquoise module are involved in both humoral immune and acute inflammatory response, and also complement activation signals [Table S3].

**Figure 3.**
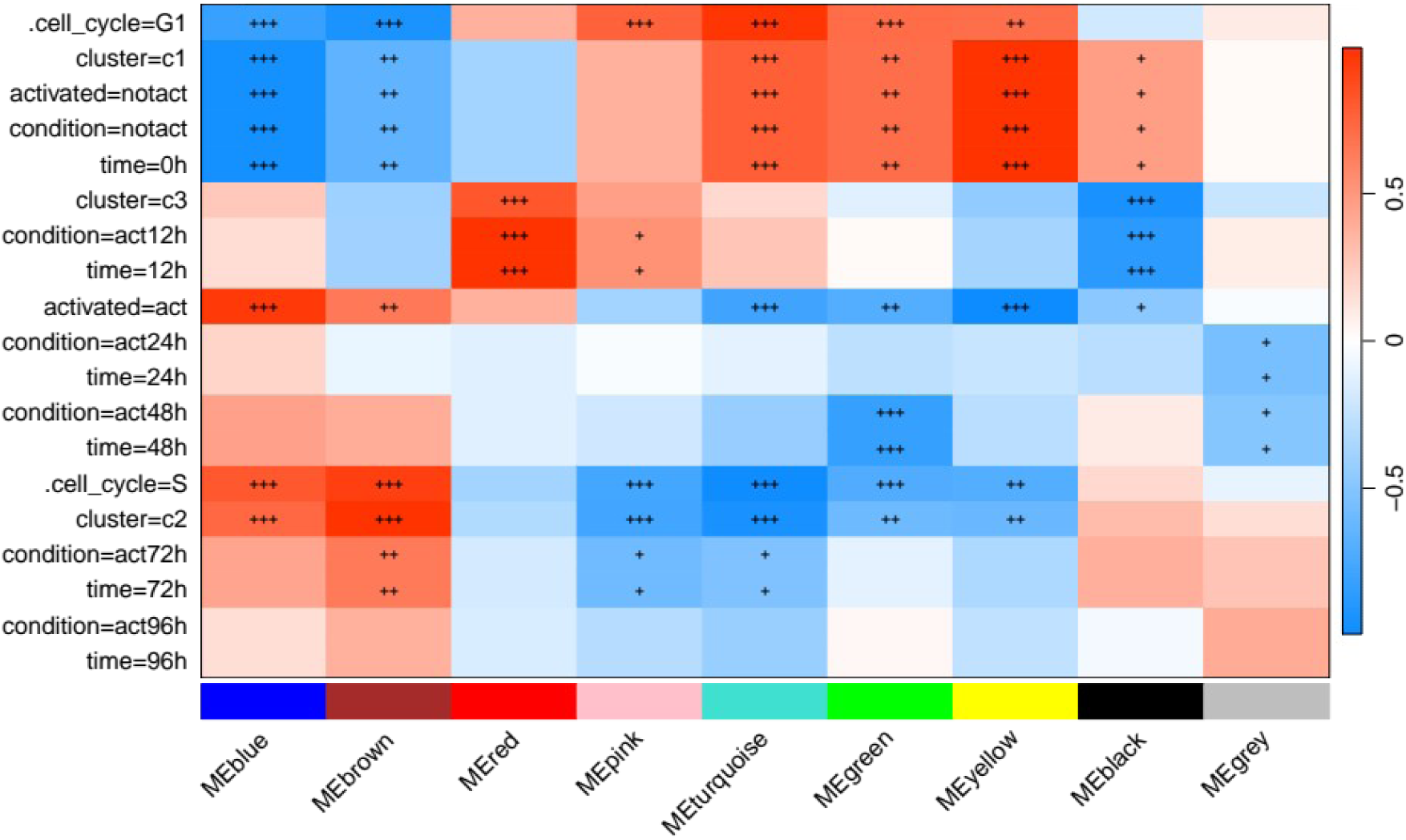
**WGCNA+ analysis of a single-omics (proteomics) dataset [**Wolf et al., *Nat Immunol* 2020]. Figure generated using the WGCNA+ *plotModuleTraitHeatmap()* function. The figure shows a heatmap of Pearson correlations between each module eigengene and each available phenotype. Red and blue colors indicate positive and negative correlation, respectively. Each cell in the heatmap is marked with one (p≤ 0.05), two (p≤ 0.01) or three (p≤ 0.001) asterisks. Cells without asterisks are p > 0.05.

Acquiring knowledge on the modules that are most associated with phenotype is central to WGCNA+. Yet, biologists often need comprehensive views of the biological roles played by the top phenotype-correlated modules. To address this, we leveraged the WGCNA+ built-in LLM-based annotation functions. For each module, these functions integrate its top correlated genes and enriched gene sets into a structured prompt which is then submitted to a LLM model to generate a grounded description of the module’s potential biological roles (Methods). LLM-aided interpretation indicated that the blue module (Fig. 4A), positively correlated with immune cell activation, evoking proliferative and biosynthetic programs. Consistent with Table S2’s enrichment for interleukin-related gene sets, top blue-module features center on MYC, E2F family cell cycle regulators, and IL-2 signaling. In activated T cells, TCR and IL-2 signals sustain Myc expression, driving metabolic reprogramming, amino-acid uptake, and bioenergetic/biosynthetic changes required for clonal expansion and effector differentiation (Preston et al., *EMBO J* 2015; Wang et al., *Immunity* 2011; Marchingo et al., *eLife* 2020). E2F factors then support the G1/S transition and DNA replication. Additionally, an enrichment for drug-related pathways and estradiol-related signals emerge (Table S2). While a mechanistic link to MYC/E2F biology is plausible (e.g., via estrogen-responsive proliferative signaling in lymphocytes), this connection is not directly established by the current data and could be considered for follow-up analyses. Collectively, these patterns are consistent with a module that captures activation-linked growth, cell cycle progression, and anabolic remodeling in T cells [Fig.4A; Table S2].

**Figure 4.**
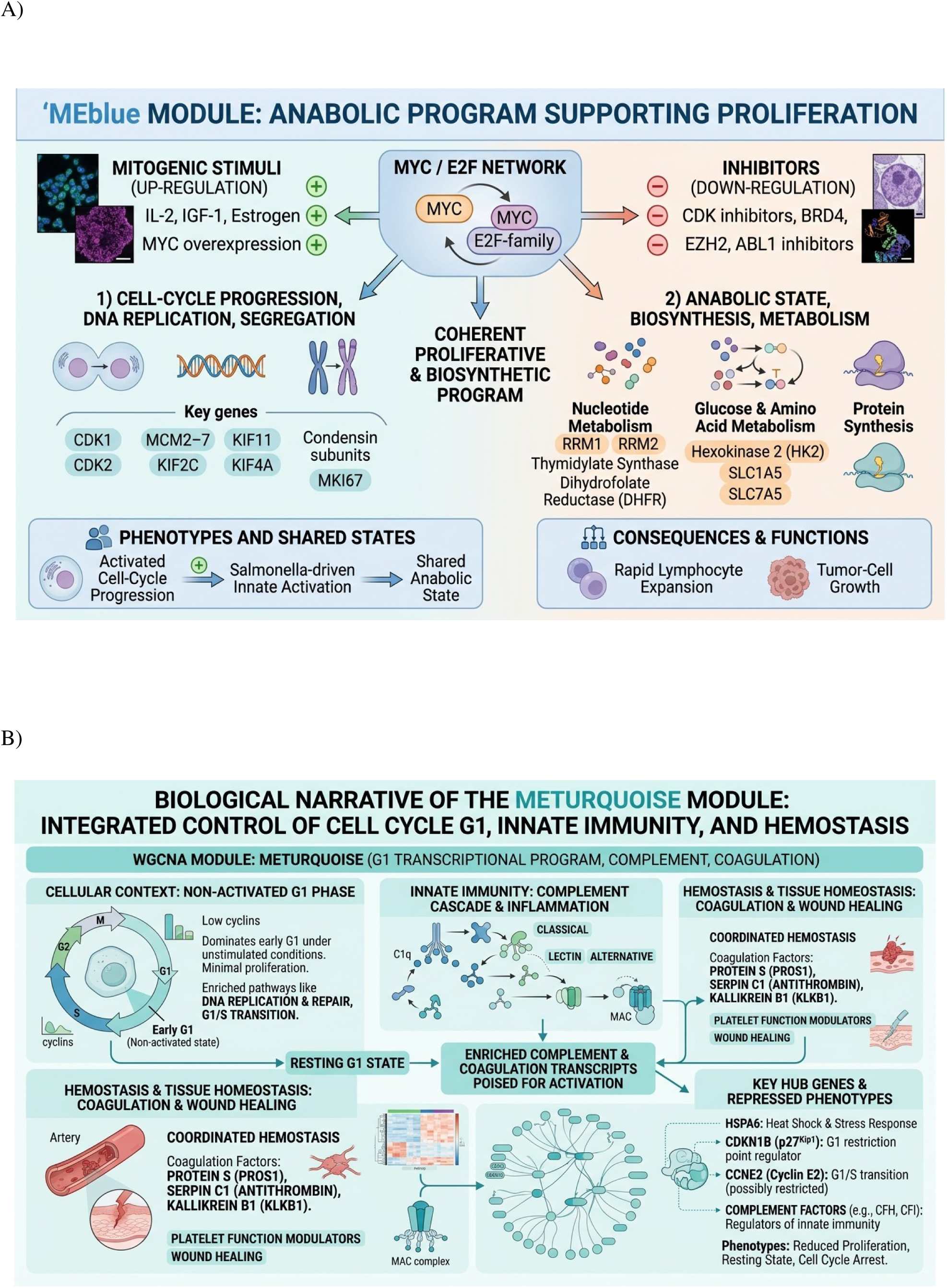
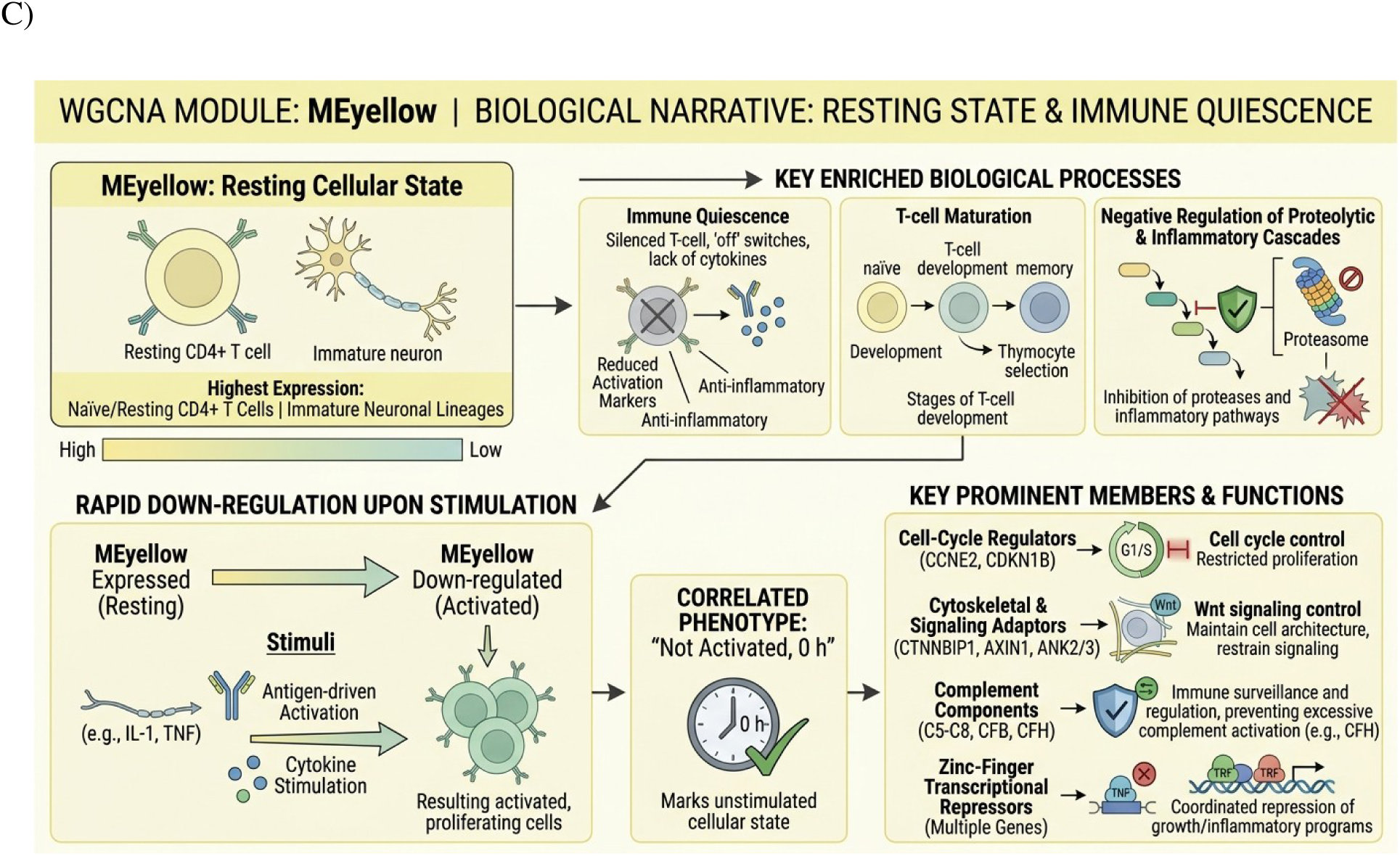
WGCNA+ integrated LLM-generated infographics produced for a single-omics (proteomics) dataset. [Wolf et al., *Nat Immunol* 2020]. The WGCNA+ modules, blue, turquoise and yellow, are interpreted by the “*gemini-3.1-flash-image-preview*” AI model. Module-specific infographics generated by the WGCNA+::*create_module_infographic()* function on blue (A), turquoise (B), and yellow module (C).

Beyond supporting current knowledge on T cell activation biology, WGCNA+ LLM indicated that the turquoise module, which negatively correlated with immune activation, relates to humoral immune and acute inflammatory response signals, including complement activation [Fig.4B]. These data are consistent with data in Table S3. Naive and quiescent T cells retain an intracellular complement system, including C3 and CD46, that sustains mTOR-dependent homeostatic survival signaling at baseline (Liszewski et al., *Immunity* 2013; West et al., *Annual Rev Immunol* 2018; Cardone et al., *Nat Immunol* 2010). The negative correlation of the turquoise module with activation in this data might reflect a decrease in baseline receptor/pathway component abundance rather than a simple on/off resting-state switch, a distinction that deserves follow-up investigations. Interestingly, quiescent T cells may enforce cell cycle arrest through inhibitors such as p27/CDKN1B, which is degraded via autophagy-dependent mechanisms upon TCR stimulation to permit G1/S entry (Jia et al., *Autophagy* 2015; Lewis et al, *Front Cell Dev Biol* 2021). The co-occurrence of complement- and cell-cycle-related features in a module that decreases with activation may be consistent with a coordinated resting-state maintenance program.

Finally, WGCNA+ also pointed to regulation of T-cell activity by the yellow module, including resting-state maintenance, T-cell maturation, and response to activating stimuli, also negatively correlated with activation [Fig.3; Fig.4C]. This is in line with the broader concept that T cell quiescence is not a passive state but is actively enforced by dedicated transcriptional and signalling programs that must be controlled for activation to proceed (Bennett et al., *Int J Mol Sci* 2020; Lewis et al, *Front Cell Dev Biol* 2021). In this context, the yellow module might reflect homeostatic circuitries that sets activation thresholds and preserves readiness in the naive/resting state. Possibly, IGF-1/IL-2-driven, TCR-independent homeostatic proliferation of Tregs cells via PI3K/Akt–STAT5 [Shapiro et al. *J Immunol* 2023] may be mechanistically more relevant to a resting/homeostatic program compared to the TCR-driven clonal expansion captured by the blue module. Altogether, these data align with results in original paper [Wolf et al, *Nat Immunol* 2020] and offer novel biological insights that deserve further studies.

WGCNA+ offers functionalities dedicated to consensus analysis. Analysis of consensus aims to define co-expression structures reproducible across distinct biological conditions (Methods). We performed consensus co-expression network analysis on liver transcriptomic data from two independent sample groups defined by sex, comprising 134 female and 125 male mice. This is a publicly-available dataset commonly used in WGCNA [https://github.com/edo98811/WGCNA_official_documentation]. First, we inferred co-expression networks separately for each sex, after which a joint consensus network was derived by requiring topological overlap to be reproducible across both sexes. WGCNA+ consensus analysis detected 13 gene modules, ranging in size from 44 (salmon) to 294 genes (turquoise), together covering 1,700 of the 2,000 analyzed genes. The remaining 300 genes were assigned to the grey module. We sought to assess how faithfully these consensus modules would reflect the structure in each sex individually. We cross-tabulated consensus module membership against the modules independently detected within female-only and male-only networks [Fig.5]. Cross-tabulations show substantial diagonal correspondence, indicating that a large fraction of features assigned to a given consensus module fall into a single matching module in each sex-specific network. We observed modest off-diagonal events into neighboring modules. We tested statistically this correspondence using Fisher’s exact test, and found that modules sharing the same color label across female and male networks exhibit significant feature-membership overlap, with FDR values from <1e-66 (purple module) to <1e-216 (turquoise module).

**Figure 5.**
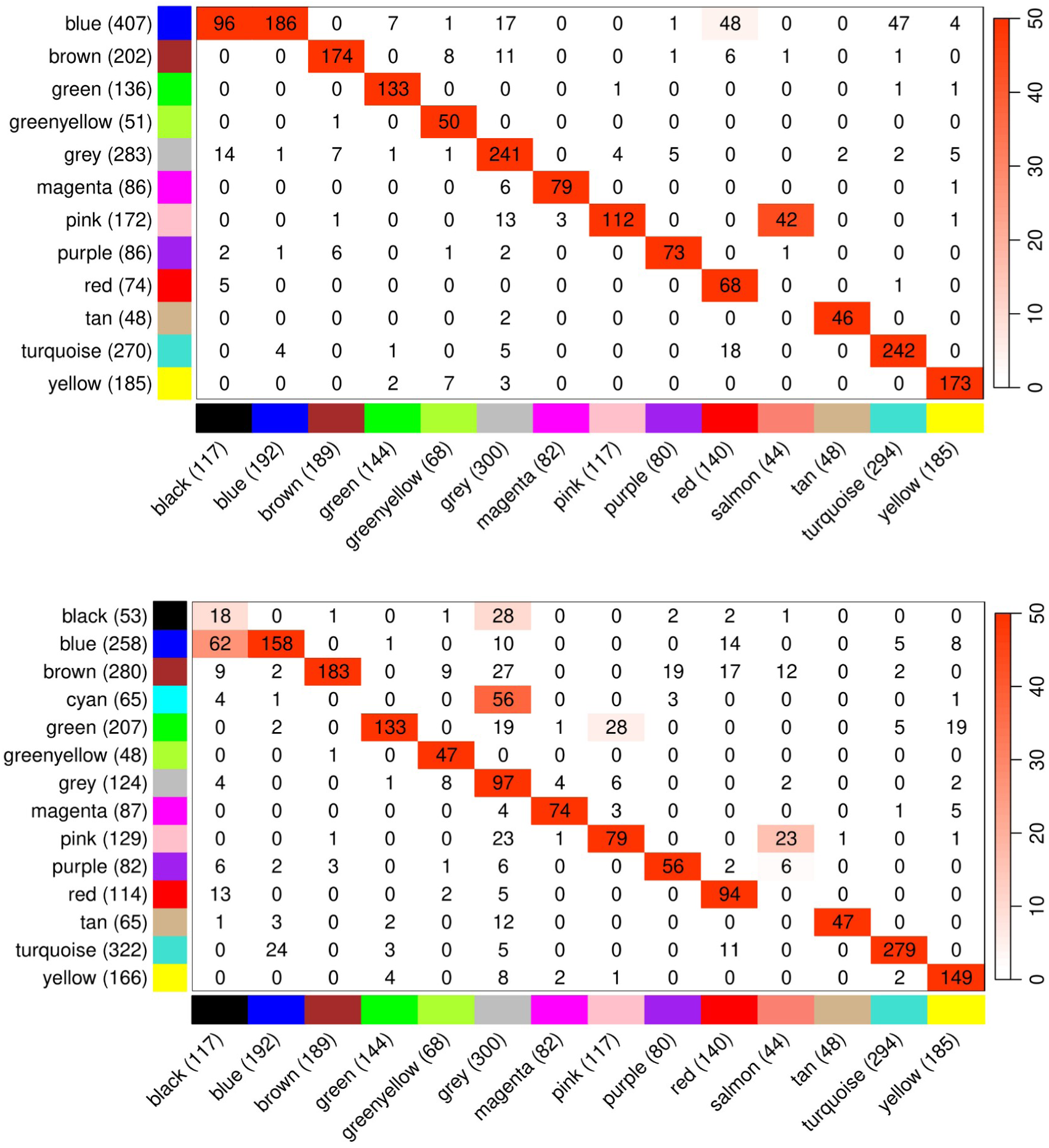
WGCNA+ consensus analysis between female and male liver transcriptomics data. WGCNA+ consensus analysis was run using the *runConsensusWGCNA()* function. Correspondence between consensus module assignments (x-axis in both maps) and the modules independently detected within the male-only (map above) and female-only (map below) liver co-expression networks. Cell values indicate the number of shared genes; the diagonal pattern shows that consensus modules closely match sex-specific modules of the same color.

Altogether, these data indicate a degree of conservation in the overall modular architecture of the liver transcriptome between sexes at this level of resolution. The largest consensus module, turquoise (294 genes), showed the strongest cross-sex reproducibility of all modules tested, suggesting it captures a core, sex-independent co-expression program rather than sex-specific signals. We next sought to gain knowledge on the consensus between the two sexes in the biological relevance of each module. We calculated the correlation between modules and measured physiological traits (length, abdominal fat, non-abdominal fat, total_fat). For each consensus module, eigengene-trait correlations were computed separately within the female and male datasets, and a module-trait relationship was retained if statistically significant (p <= 0.05) and concordant in sign within both sexes independently [Fig.6]. Consistent consensus signals involved adiposity-related traits. The pink co-expression module exhibited a strong negative consensus correlation with both non-abdominal and total-fat mass. On the other hand, the black, red, and blue co-expression modules each showed positive consensus correlations with adiposity traits [Fig.6]. Other co-expression modules, despite structural reproducibility, showed no trait association and sign consistent across both sexes. This might be explained by lack of relationship to the measured physiological traits, or a certain degree of sex-dependency.

**Figure 6.**
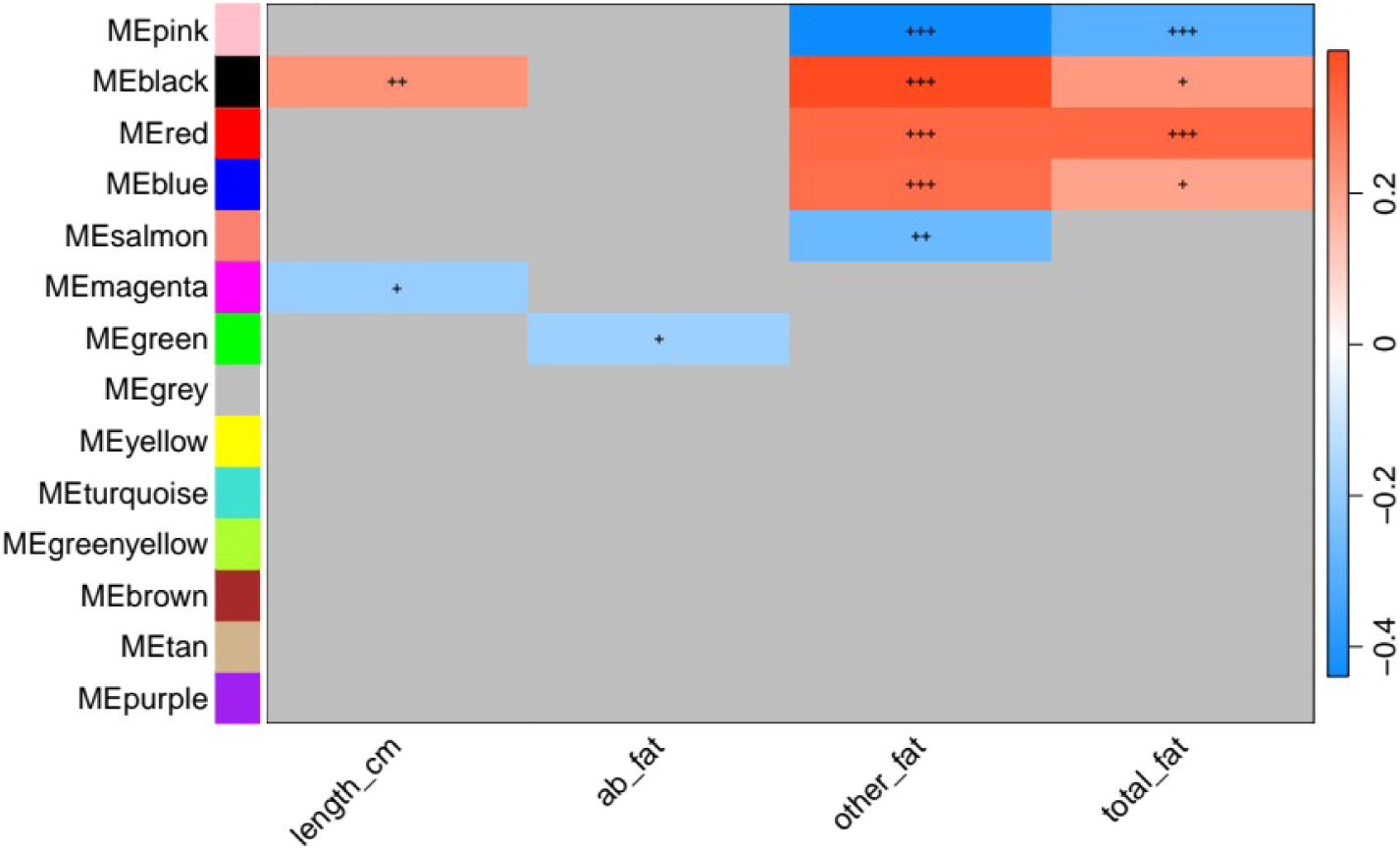
Consensus module-trait relationships. Heatmap generated using WGCNA+ *plotLabeledCorrelationHeatmap()*. Cells are colored by correlation strength (red = positive, blue = negative) and shown only where the module-trait association was statistically significant (p <= 0.05) and sign-concordant in both female and male networks independently; grey cells indicate no significant consensus relationship. Significance is indicated by “+” symbols (+ p<0.05, ++ p<0.01, +++ p<0.001).

While the consensus analysis identified co-expression modules concordant across the sexes, a complementary question concerns whether modules are also structurally preserved. Preservation would point to genes that co-vary in one sex, while also maintaining similar connectivity patterns and network topology in the other sex. Beyond consensus, a central question in multi-omics co-expression analysis is whether gene modules are maintained across biological layers. The possibility that genes that co-express at the transcript level also co-vary at the protein, metabolite or miRNA level would have high biological impact. Co-expression modules preserved across data types or phenotypes would represent functionally coherent regulatory programs rather than layer-specific signals. For preservation, a test layer is tested against a ‘reference’ layer (Methods). We conducted WGCNA+ module preservation analysis in the liver expression dataset to test preservation in liver transcriptomics between the sexes. The liver is a sexually dimorphic organ in mammals, especially structurally and metabolically [Matz-Soja et al., *Journal of Hepatology* 2026]. Previous studies have reported differential expression profiles between the sexes in the liver [Delongchamp et al, *BMC Bioinformatics* 2005; Rajcsanyi et al, *Scientific Reports* 2025]. It remains an intriguing possibility that core transcriptomic networks share common regulatory programs in the male and female liver. We defined modules in the female dataset as the reference, and tested for structural preservation in the male dataset using WGCNA’s Z-summary statistic, which integrates multiple density and connectivity measures. We found 5 modules (green, blue, turquoise, brown, cyan) to be strongly preserved, 3 modules (royalblue, lightcyan, greenyellow) moderately preserved, and 2 modules (lightyellow, pink) to exhibit lower preservation [Fig. 7A-B]. Altogether, these data indicate substantial structural preservation in liver transcriptomic profiles, suggesting sharing in core co-expression architecture across the sexes. In parallel, weakly preserved modules might indicate sex-specific programs. To highlight modules that are both significantly preserved and trait-correlated, WGCNA+ allows to weight the Z-score by geometric mean of the module-trait correlation across datasets [Fig. 7B].

**Figure 7.**
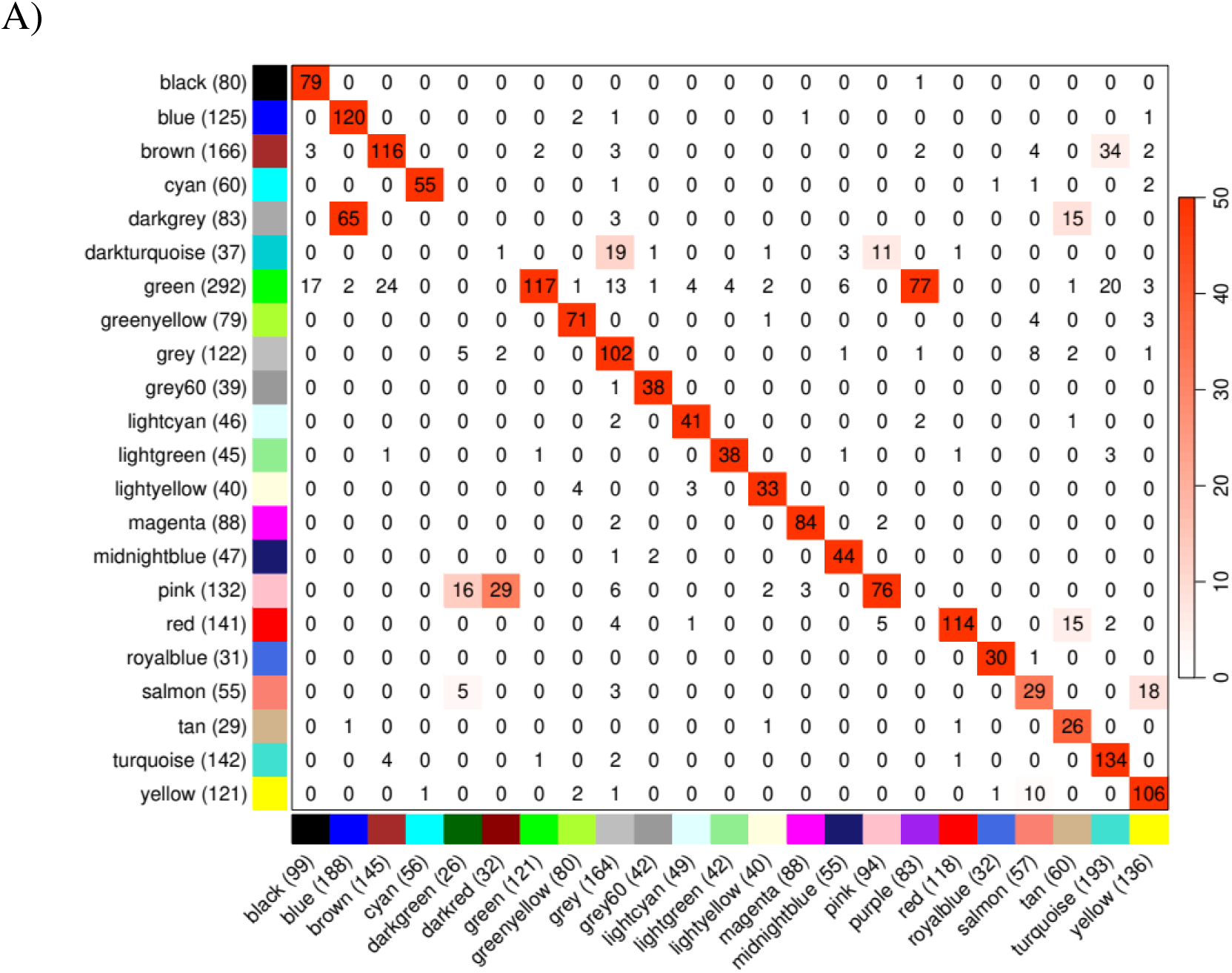
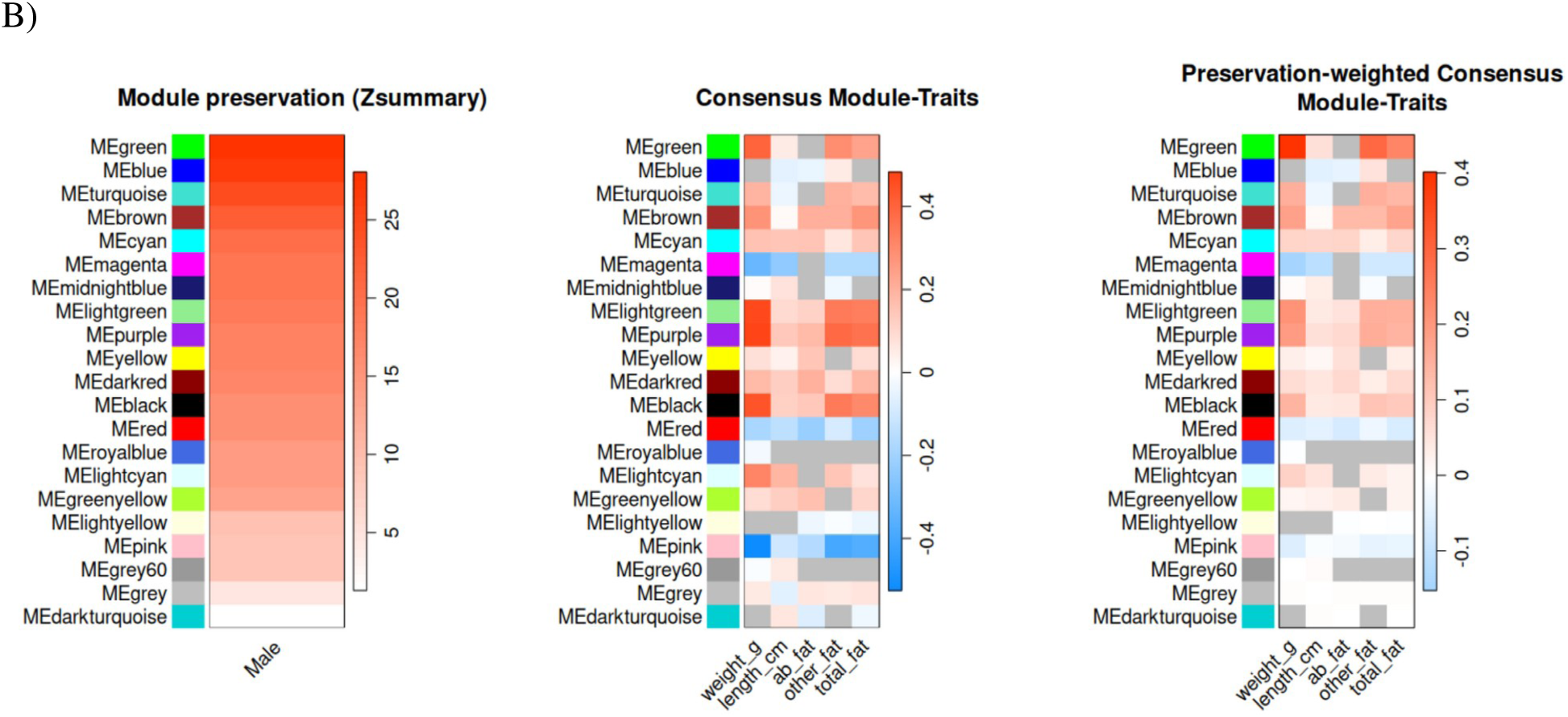
WGCNA+ preservation analysis (tested using *runPreservationWGCNA()*) of male and female mouse liver transcriptomics [https://github.com/edo98811/WGCNA_official_documentation]. A) Fig. generated with *plotConsensusOverlapHeatmap()*. The heatmap provides data on module correspondence between the male network and the female-male consensus network. On the y-axis, the number of features mapped in the module in shown in parentheses. The number of features shared between the module pair is reported within each cell. The cell color corresponds to the Fisher’s exact test p-value (-log10 scale; white to red: low to highly significant overlap). B) Fig. generated with *plotPreservationModuleTraits()*. On the left, heatmap of Z-summary values per module. Higher values indicate higher preservation. In the middle, heatmap of consensus module trait, plotting geometric mean consensus of module-trait correlations. On the right, heatmap of weighted preservation: the mean of squared Z-score is calculated per module across all datasets and divided by the maximum across all modules to normalize into [0,1]. Thus, the output values are a score of module’s preservation strength. This value is then multiplied by the geometric mean of module-trait correlation across datasets.

To get insights on the biological relevance of top preserved modules, we performed gene set and pathway enrichment analysis of the green, blue, turquoise and brown modules, using gprofiler2 [Kolberg et al., *F1000res* 2020]. We tested enrichment for ‘GO:BP’, ‘GO:MF’, ‘KEGG’ and ‘REAC’ using the *gprofiler2::gost()* function. We selected terms with p<0.01 and a minimum overlap with the target gene set or pathway of 20 features. We found enrichment for metabolic process and biosynthesis terms, immune regulation terms, and varied cellular control processes [Table S4]. Rather than a single shared signature, each module carries a distinct, largely non-overlapping biological theme, suggesting that preserved co-expression structure partitions liver homeostatic biology into separable functional programs.

The blue module is enriched for broad metabolic-regulatory terms, including “regulation of primary metabolic process” (GO:0080090), “regulation of metabolic process” (GO:0019222), “regulation of biosynthetic process” (GO:0009889), and their negative counterparts (”negative regulation of metabolic process,” GO:0009892; “negative regulation of biosynthetic process,” GO:0009890). As none of these terms appear among the top hits for brown, turquoise, or green, this metabolic signature appears as blue-module-specific feature. The blue module additionally shows strong enrichment for gene expression and RNA metabolism control, including “regulation of gene expression” (GO:0010468), “regulation of RNA metabolic process” (GO:0051252), “regulation of transcription by RNA polymerase II” (GO:0006357), and negative regulators of transcription (GO:0045892, GO:0000122). These data suggest the blue module encodes transcription factors, chromatin regulators and RNA-processing machinery that tune hepatocyte metabolic gene programs, consistent with the WGCNA-based liver homeostasis literature (Esmaili et al., *Cell Syst* 2021; Hu et al., *Genom Inform* 2023).

The brown and turquoise modules are dominated almost exclusively by immune and defense-related terms rather than metabolic terms. Brown shows top enrichment for “immune system process” (GO:0002376), “defense response” (GO:0006952), “immune response” (GO:0006955), “innate immune response” (GO:0045087), and cytokine production (GO:0001816/7), together with KEGG pathways such as Fc gamma R-mediated phagocytosis, Phagosome, Toll-like receptor signaling, and REACTOME “Innate Immune System” pathway. Turquoise shows an even larger immune signature, including immune system process and its regulation, defense response, leukocyte/T-cell/lymphocyte activation and proliferation, cytokine-mediated signaling (GO:0019221), and adaptive immune response (GO:0002250), extending brown’s innate-immunity theme into adaptive and lymphocyte-specific processes.

The green module is enriched for a third, distinct theme: system and tissue development (GO:0048731), vasculature and blood vessel development (GO:0001944, GO:0001568), extracellular matrix organization (GO:0030198), together with cell-surface receptor signaling (GO:0007166) and regulation of signal transduction (GO:0009966). This is reinforced by cellular component and pathway terms, including extracellular matrix, and KEGG/REACTOME pathways including Integrin signaling, Focal adhesion, PI3K-Akt signaling, ECM-receptor interaction, and Collagen formation/biosynthesis. This module reflects signaling and developmental programs that establish and maintain liver architecture, vascularization and hepatocyte identity, rather than intrinsic metabolic flux control.

Altogether, these data suggest that preserved networks across the sexes are involved in constitutive physiological processes. The four preserved modules partition into complementary biological programs: blue governs core metabolic and transcriptional regulation, brown and turquoise govern innate and adaptive immune surveillance, and green governs structural, vascular and developmental signaling. Their joint preservation across male and female liver transcriptomes suggests that these module-specific regulatory circuits are each independently conserved, key components of hepatic homeostasis in both male and female liver (Esmaili et al., *Cell Syst* 2021; Hu et al., *Genom Inform* 2023).

WGCNA+ offers unique functionalities for multi-omics data. We used a multi-omics breast cancer dataset downloaded from the TCGA repository [https://portal.gdc.cancer.gov/]. The dataset, herein referred to as ‘mox-brca’, comprises 150 samples concurrently profiled in transcriptome (‘gx’), proteome (‘px’), and miRNAs (‘mir’). The ‘mox-brca’ offers to biologists an excellent opportunity to assess the power of WGCNA+ in a large, real-world biological data and cross check biologically meaningful findings against the extensive literature for breast cancer. Multi-omics data are typically generated by distinct technologies, each with its own detection limit, and exhibit different statistical distributions and noise profiles. Distinct data types often require tailored parametrization in bioinformatic analyses. WGCNA+ initially performs hierarchical clustering for each data type independently [Fig.8A], allowing evaluation of key parameters -primarily power- for each datatype. Assessment of module-traits association patterns indicated that the WGCNA+ gx blue module is significantly associated with basal breast cancer samples, while the px green module is significantly associated with Her2 positive breast cancer samples [Fig.8B]. By inspecting the multi-omics features mapped into the green module, we found that HER2, HER2_pY1248, HER3_pY1289 and EGFR are components of the module. This finding, in fact, provides the rationale for the exclusive association between the green module and Her2-positive breast cancer. The detection of the main HER2 protein isoform and its variants strongly supports the biological relevance of the WGCNA+-inferred modules to the phenotype.

**Figure 8.**
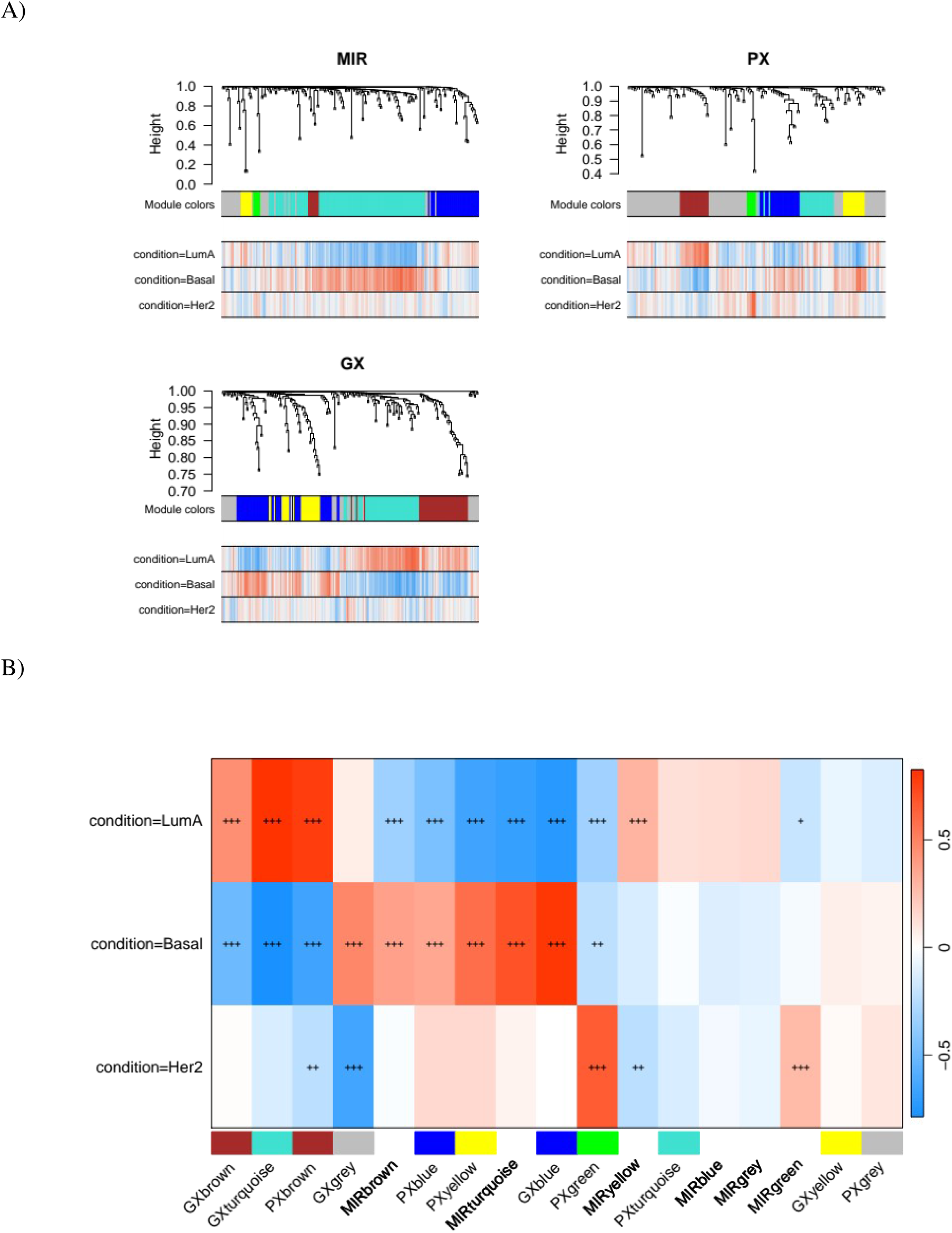
WGCNA+ analysis of multi-omics data. A) Fig. generated using WGCNA+ *plotMultiDendroAndColors()*. Dendrogram of the features’ hierarchical clustering tree built from consensus TOM per data type. The figure enables to assess consistency in module structures between data types. B) Fig. generated using WGCNA+ *plotModuleTraitHeatmap()*. Heatmap of Pearson correlations between module eigengenes and phenotypes. Red and blue indicate positive and negative correlation, respectively. Each heatmap cell is marked with one (p≤ 0.05), two (p≤ 0.01) or three (p≤ 0.001) asterisks. Cells without asterisks are p > 0.05.

We sought to acquire a comprehensive overview of all possible relationships between module eigengenes and phenotypes across all omics layers. We reasoned that this unbiased, systems-level approach would allow us to identify biologically coherent, cross-omics associations with phenotype that might otherwise be missed by hypothesis-driven or single-layer analyses. To this end, we first computed the correlation between each module eigengenes and each available trait, within each data type. To identify the top modules within each layer, we scored each module by its overall trait association strength and selected the modules scoring within 90% of the best-scoring module in that layer. The use of this layer-specific relative threshold, rather than absolute or fixed cutoff, enabled us to identify top modules contextualized to the experiment. We observed both clusters driven by negative correlation, and clusters driven by positive correlation [Fig.9]. For instance, we detected a general positive correlation between the gene expression modules brown, yellow and red, miRNA modules turquoise, green, and yellow, and proteomics module green, suggestive of potential cross-omics regulation [Fig.9].

**Figure 9.**
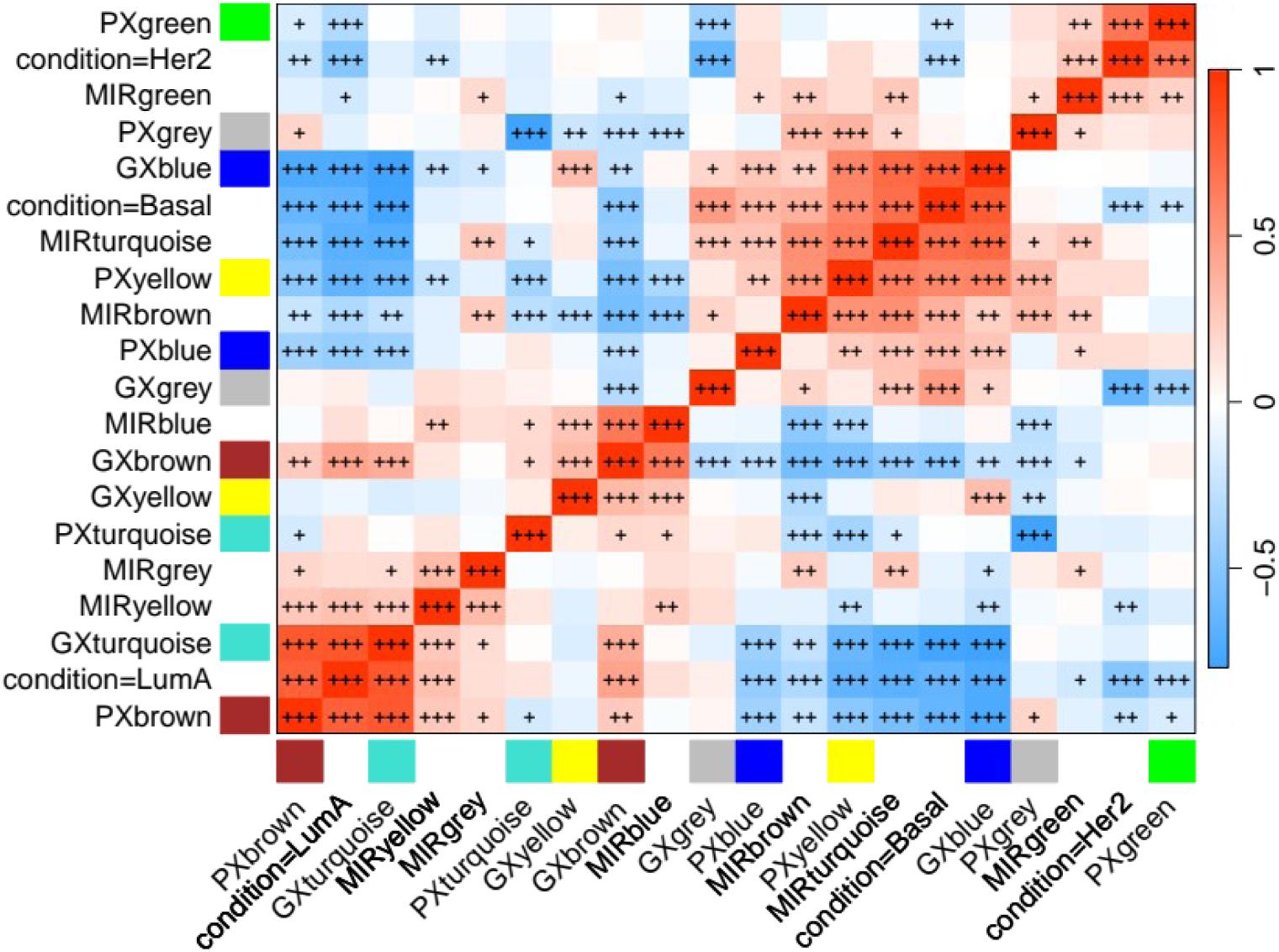
WGCNA+ analysis of the multi-omics dataset mox-brca. Figure generated using WGCNA+ *plotEigenGeneAdjacencyHeatmap()*. The figure shows a heatmap of Pearson correlations between all combinations of module eigengenes and available phenotypes. Red and blue color indicate positive and negative correlation, respectively. Each cell in the heatmap is marked with one (p≤ 0.05), two (p≤ 0.01) or three (p≤ 0.001) asterisks. Cells without asterisks are p > 0.05.

Previous analyses provided data about module-level cross-omics relationships. However, the degree to which individual features are involved into inter-layer relationship remains uncharacterized. To address this question, we applied the LASAGNA multi-layer model (Methods) to the mox-brca dataset, constructing a layered graph in which transcriptomics, proteomics, and miRNA features are represented as distinct node partitions connected by phenotype-weighted inter-layer edges [Fig.10]. Conditioning the graph against the HER2 phenotype (Methods) revealed a tightly connected inter-layer subnetwork, linking HER2 receptor protein nodes in the proteomics layer to CDK18 and NDRG2 transcriptomic features. Intriguingly, a set of miRNAs is also detected both with positive and anti-correlation patterns, including mir-455 and mir-20a [Fig.10]. This feature-resolution view complements the module-level analysis and may pinpoint specific cross-omics regulatory relationships that analyses at the module level alone, including module eigengenes, could not reveal.

**Figure 10.**
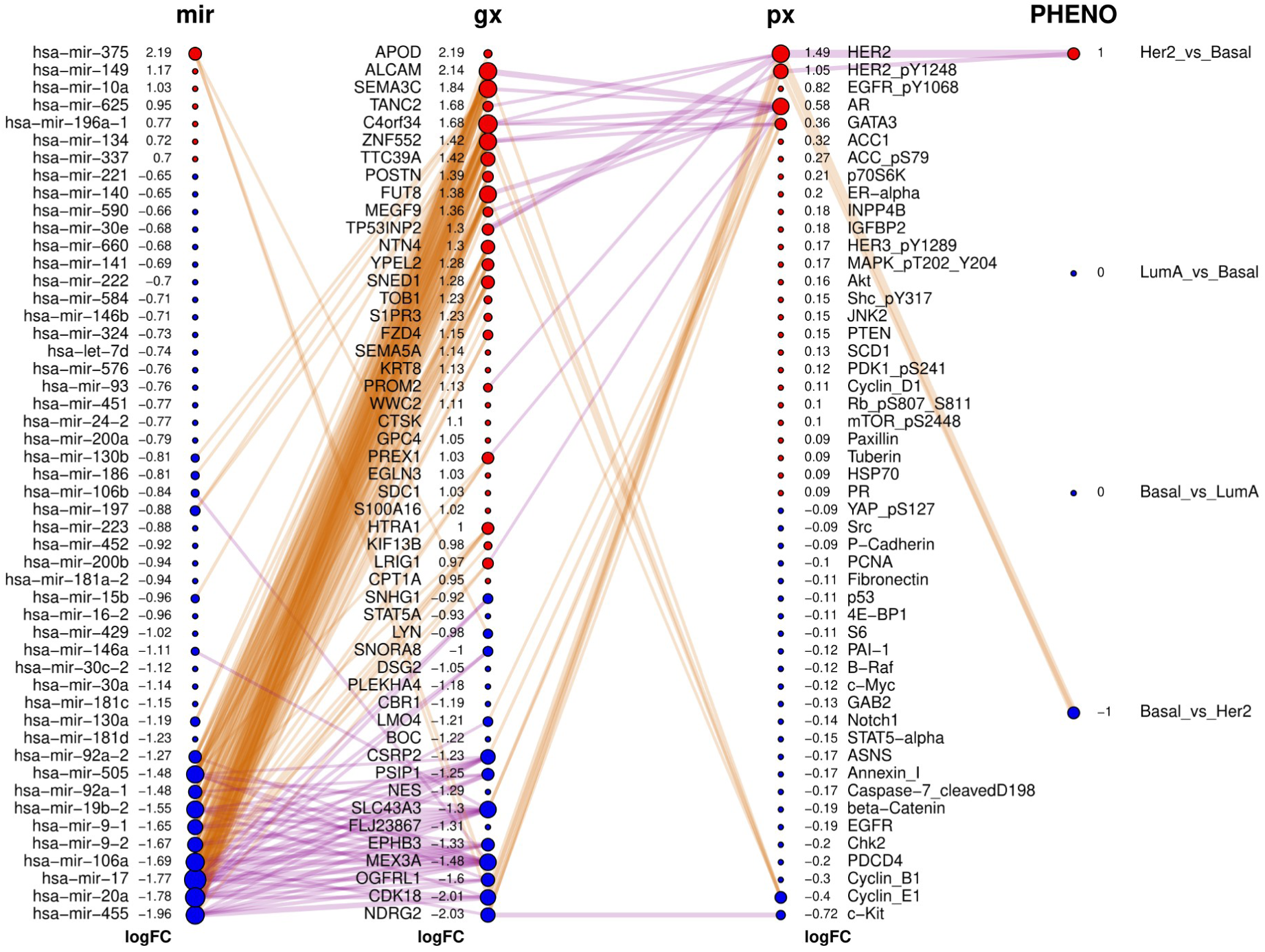
LASAGNA feature-level multi-partite graph. Figure generated using LASAGNA *plot_multipartite()*. Each layer correspond to a data type (mir=miRNA; gx=transcriptomics; px=proteomics). Nodes in each layer are features. Features are ordered by Log2FC for the contrast of interest (Basal_vs_Her2). Red and blue nodes indicate positive and negative Log2FC, respectively. Edges between layers represent correlation between features. Purple and yellow edges indicate positive and negative correlation, respectively. Thicker edges represent higher correlation coefficient. The sizes of circles represent the feature’s page-rank centrality.

## 4. Discussion

The WGCNA R package provides functionalities to identify correlated genes modules and their correlation with phenotypes [Langfelder & Horvath, *BMC Bioinformatics* 2008]. However, it provides no mechanism to jointly model heterogeneous omics layers with different feature space and scales. Furthermore, it does not provide functionalities to connect genes with other omics layers, such as proteins, metabolites, lipids etc. There are functionalities, such as those for consensus WGCNA, capable of handling multi-data structures but these largely refer to multiple datasets of the same molecular type rather than multi-omics data. There is a pressing need to extend current WGCNA to cover cross-layers relationships and be directly applicable to multi-omics data.

To address this problem, we have developed WGCNA+, the first framework, to our knowledge, that delivers an end-to-end, multi-omics extension of WGCNA. WGCNA+ combines per-layer inference, cross-layer enrichment, consensus/preservation testing, and computational acceleration within a single, WGCNA-compatible package. WGCNA+ substantially enhances the standard WGCNA framework by enabling integrated, joint analysis of multiple omics layers, including but not limited to transcriptomics, proteomics and metabolomics. We have demonstrated WGCNA+ capabilities in identifying biologically meaningful signals in single and multi-omics data, spanning diverse scenarios in sample size, omics layers and phenotypic breadth. By identifying co-expression modules robustly and concurrently altered across biological layers, and significantly associated with phenotypes, WGCNA+ directly empowers researchers to detect biological processes characterized by system-wide molecular alterations.

WGCNA+ also offers dedicated approaches to test preservation of multi-omics modules, enabling evaluation of unique or shared biological alterations across conditions. This functionality is critical for distinguishing conserved mechanisms from phenotype-specific signatures, and would serve existing personalized medicine studies well. Inferring structural preservation has important implications beyond agreement in eigengene direction or trait association. A module may be concordant yet structurally reorganized, reflecting rewired regulatory architecture rather than conserved co-expression program.

Our work also comprises the release of a standalone, innovative visualization tool of multi-dimensional data, LASAGNA (Layered Approach for Simultaneous Analysis of Genomic and Network Associations). The LASAGNA model provides a multi-condition network representation that links per-layer profiling and phenotypic outcomes through a structured graphical model. In a complex multi-omics breast cancer dataset, we demonstrate the LASAGNA’s capabilities of showing relationship between multiple omics data, while incorporating effect size of differential expression between phenotypes, effect direction, cross-omics feature correlation, and correlation with phenotypes. Modern biomedical research is increasingly relying on multi-omics profiling to resolve complex diseases [Hasin et al, *Genome Biology* 2017]. Unlike conventional network visualization tools, which typically operate on a single correlation matrix agnostic to phenotype, LASAGNA provides an innovative multi-modal framework that combines phenotype-conditioned edge weighting and feature-resolution cross-omics graphs. Because LASAGNA operates on any correlation-derived multi-omics matrix, its relevance extends beyond the WGCNA+ framework and it can serve as a general-purpose tool for the wide multi-omics community.

Taken together, WGCNA+ and LASAGNA jointly redefine a co-expression analysis toolkit. WGCNA+ establishes the statistical and computational backbone for multi-omics module discovery, while the LASAGNA model supplies the phenotype-aware visualizations with feature-resolved graphs. This combination contributes toward closing a gap that has long persisted in bioinformatics, and may position the WGCNA+/LASAGNA framework as a new standard for multi-omics systems biology.

WGCNA+ uniquely incorporates functionalities for automated LLM-driven result interpretation. To our knowledge, no other WGCNA-derived or network-based multi-omics tool integrates automated LLM-driven interpretation and infographic synthesis directly into the analysis pipeline. This advance makes WGCNA+ a self-documenting discovery platform rather than a purely computational one, as it effectively provides scientists with immediate descriptions of the identified modules and their potential biological relevance. Our benchmarking analyses demonstrate that the WGCNA+ AI-reports are cross-disciplinary in nature, remain focused on the investigated phenotype, and show reasonable consistency across iterations. This capability dramatically accelerates evaluation, reporting, and decision-making processes, enhancing the progression toward validatory functional genomics screens.

Unfortunately, pharmaceutical R&D pipelines remain very time-consuming and in many cases, inefficient. Partly, this inefficiency stems from the absence of tools capable of (i) performing integration of heterogeneous omics layers, (ii) infer druggable, phenotype-associated alterations from intricate molecular networks and (iii) offering scientists with an immediate, interpretable summary of the findings. Bridging these gaps is essential to support modern biomedical research. WGCNA+ offers concrete steps toward directly addressing these critical needs. The integrative capabilities in WGCNA+, comprising multi-omics data integration and AI-powered interpretation of results, move beyond single-layer correlations and toward a genuinely systems-level understanding of phenotype biology. Altogether, WGCNA+ represents a step toward more efficient, data-driven translational research.

## Acknowledgements

We thank the BigOmics Analytics team for feedback on this work.

